# Chromosome-level genome assembly of a human fungal pathogen reveals synteny among geographically distinct species

**DOI:** 10.1101/2021.07.13.452254

**Authors:** Mark Voorhies, Shirli Cohen, Terrance P. Shea, Semar Petrus, José F. Muñoz, Shane Poplawski, William E. Goldman, Todd P. Michael, Christina A. Cuomo, Anita Sil, Sinem Beyhan

**Author notes:** These authors contributed equally to this work (Mark Voorhies, Shirli Cohen). Plant Molecular and Cellular Biology Laboratory, The Salk Institute for Biological Studies, La Jolla, CA 92037. Co-corresponding authors: Sinem Beyhan J. Craig Venter Institute, 4120 Capricorn Lane, La Jolla, CA 92037, Anita Sil Department of Microbiology and Immunology, University of California San Francisco, San Francisco, CA 94143-0414, Christina A. Cuomo Broad Institute of MIT and Harvard, 415 Main Street, Cambridge, MA 02142.

## Abstract

*Histoplasma capsulatum*, a dimorphic fungal pathogen, is the most common cause of fungal respiratory infections in immunocompetent hosts. *Histoplasma* is endemic in the Ohio and Mississippi River Valleys in the United States and also distributed worldwide. Previous studies revealed at least eight clades, each specific to a geographic location: North American classes 1 and 2 (NAm 1 and NAm 2), Latin American groups A and B (LAm A and LAm B), Eurasian, Netherlands, Australian and African, and an additional distinct lineage (H81) comprised of Panamanian isolates. Previously assembled *Histoplasma* genomes are highly fragmented, with the highly repetitive G217B (NAm 2) strain, which has been used for most whole genome-scale transcriptome studies, assembled into over 250 contigs. In this study, we set out to fully assemble the repeat regions and characterize the large-scale genome architecture of *Histoplasma* species. We re-sequenced five *Histoplasma* strains (WU24 (NAm 1), G217B (NAm 2), H88 (African), G186AR (Panama), and G184AR (Panama)) using Oxford Nanopore Technologies long-read sequencing technology. Here we report chromosomal-level assemblies for all five strains, which exhibit extensive synteny among the geographically distant *Histoplasma* isolates. The new assemblies revealed that *RYP2*, a major regulator of morphology and virulence, is duplicated in G186AR. In addition, we mapped previously generated transcriptome datasets onto the newly assembled chromosomes. Our analyses revealed that the expression of transposons and transposon-embedded genes are upregulated in yeast phase compared to mycelial phase in G217B and H88 strains. This study provides an important resource for fungal researchers and further highlights the importance of chromosomal-level assemblies in analyzing high-throughput datasets.

**Importance:** *Histoplasma* species are dimorphic fungi causing significant morbidity and mortality worldwide. These fungi grow as mold in the soil and as budding yeast within the human host. *Histoplasma* can be isolated from soil in diverse regions, including North America, South America, Africa and Europe. Phylogenetically distinct species of *Histoplasma* have been isolated and sequenced. However, for the commonly used strains, genome assemblies have been fragmented, leading to underutilization of genome-scale data. This study provides chromosome-level assemblies of the commonly used *Histoplasma* strains using long-read sequencing technology. Comparative analysis of these genomes shows largely conserved gene order within the chromosomes. Mapping existing transcriptome data on these new assemblies reveals clustering of transcriptionally co-regulated genes. Results of this study highlight the importance of obtaining chromosome-level assemblies in understanding the biology of human fungal pathogens.

## Introduction

*Histoplasma* and the closely related pathogens *Blastomyces*, *Coccidioides*, and *Paracoccidioides* are thermally dimorphic fungi that cause fungal infections in immunocompetent and immunocompromised hosts (1). Histoplasmosis, a systemic infection caused by *Histoplasma*, is a significant cause of mortality in immunocompromised individuals (2). Although aggressive treatment with antifungals can be successful in clearing the infection, the mortality rate is estimated to be over 50% in some regions of the world (3, 4).

*Histoplasma* is endemic in the Ohio and Mississippi River Valleys in the United States and also distributed worldwide, mostly in North America, South America and Africa. Several *Histoplasma* genomes are publicly available and have been used for detailed analysis of fungal genetics. Seminal studies by Kasuga et al. classified *Histoplasma* isolates into at least eight geographically isolated clades: North American classes 1 and 2 (NAm 1 and NAm 2), Latin American groups A and B (LAm A and LAm B), Eurasian, Netherlands, Australian and African, as well as a distinct lineage (H81) comprised of three Panamanian isolates (5, 6). Later, six phylogenetic groups within the Latin American groups were described (7). Further phylogenetic analysis of 30 additional unassembled genomes (10 NAm1, 11 NAm2, 4 LAm A, 3 Panama, and 2 Africa) divided *Histoplasma* species into five genetically distinct lineages, and four of them have been renamed as follows: *Histoplasma capsulatum* (H81 lineage), *Histoplasma mississippiense* (NAm 1), *Histoplasma ohiense* (NAm 2), and *Histoplasma suramericanum* (LAm A) (8).

*Histoplasma* exists in a saprophytic hyphal form in the soil and produces asexual spores termed conidia. Once conidia are inhaled by the mammalian host, the fungus transitions into its pathogenic yeast form, which can proliferate within host macrophages and cause disease. We and others have been studying the temperature-regulated gene networks that control cell morphology and virulence in *Histoplasma*. We have shown that four transcriptional regulators, Ryp1,2,3,4, are major regulators of cell morphology and virulence gene expression, and are required for yeast-phase growth (9–11) while the signaling mucin Msb2 is required for hyphal-phase growth (12). In these studies, we have used a wide variety of high-throughput molecular biology techniques, including transcriptional profiling, chromatin immunoprecipitation-on-chip analyses, forward genetic screens and mapping genomic modifications, to study the biology of this important fungal pathogen. However, any chromosome-level patterns in these data were obscured due to the fragmented nature of the existing *Histoplasma* genome assemblies.

Emerging sequencing technology provides complete genome assemblies that can be leveraged in high-throughput analyses. Specifically, long-read sequencing tools such as Oxford Nanopore Technologies have demonstrated significant promise in the field of fungal genomics (13–16). In this study, we re-sequenced five strains of *Histoplasma*: G217B, H88, G184AR, G186AR, and WU24, that belong to four distinct populations (5). Using a combination of long-read (Oxford Nanopore (ONT)) and short-read (Illumina) sequencing, we *de novo* assembled all five genomes to the chromosomal level. Comparison of the assembled genomes unveiled largely syntenic regions of the chromosomes. Further examination of these genomes revealed that a region containing *RYP2*, a regulator of yeast-phase growth, is duplicated in the *Histoplasma* G186AR strain. Re-analyses of high-throughput datasets also revealed that the transposon genes, as well as genes embedded in transposon-rich regions, display more abundant transcription in the yeast phase in G217B and H88 strains. This study highlights that complete genome assemblies allow new insights to be drawn from genomic data sets and will open opportunities for future studies of this organism.

## Results

### Re-sequencing of *Histoplasma* strains reveals chromosomal-level genome assemblies

Previously assembled *Histoplasma* genomes are highly fragmented, with the highly repetitive G217B strain having 261 contigs (Table S1). In order to achieve a full assembly of the repeat regions and characterize the large-scale genome architecture of *Histoplasma* species, we re-sequenced five *Histoplasma* strains (WU24, G217B, H88, G186AR, and G184AR) using Oxford Nanopore Technologies (ONT) to assemble complete chromosomes followed by polishing using Illumina short reads (Table S1 and Fig. S1). The five strains are derived from four distinct populations (5), three of which have recently been proposed as distinct species (7, 8). WU24 is from the North American 1 (NAm1) clade and renamed as *Histoplasma mississippiense*; G217B is from North American 2 (NAm2) and renamed as *Histoplasma ohiense*; G186AR and G184AR are Panamanian from H81 lineage and are renamed as *Histoplasma capsulatum*; and H88 is from the African clade (5–8).

Our results revealed that the genomes of G186AR (31 Mb), G184AR (31 Mb), and WU24 (32 Mb) are significantly smaller than H88 (38 Mb) and G217B (40 Mb), consistent with previously observed coverage differences (8). For two of the *Histoplasma* strains, WU24 and H88, we achieved complete telomere-to-telomere assemblies; H88 has 6 chromosomes while WU24 has 7 chromosomes (Fig. 1). The near-complete assemblies have 7 large contigs each, with either 12 (G186AR) or 11 (G217B and G184AR) assembled telomeres. For the G217B and G184AR assemblies, we were able to locate the 45S rDNA repeat arrays near the ends of large contigs, consistent with their subtelomeric positioning in other strains (Fig. 1). These observations are consistent with either 6 or 7 chromosomes for the G217B, G186AR and G184AR strains.

**Fig. 1.**
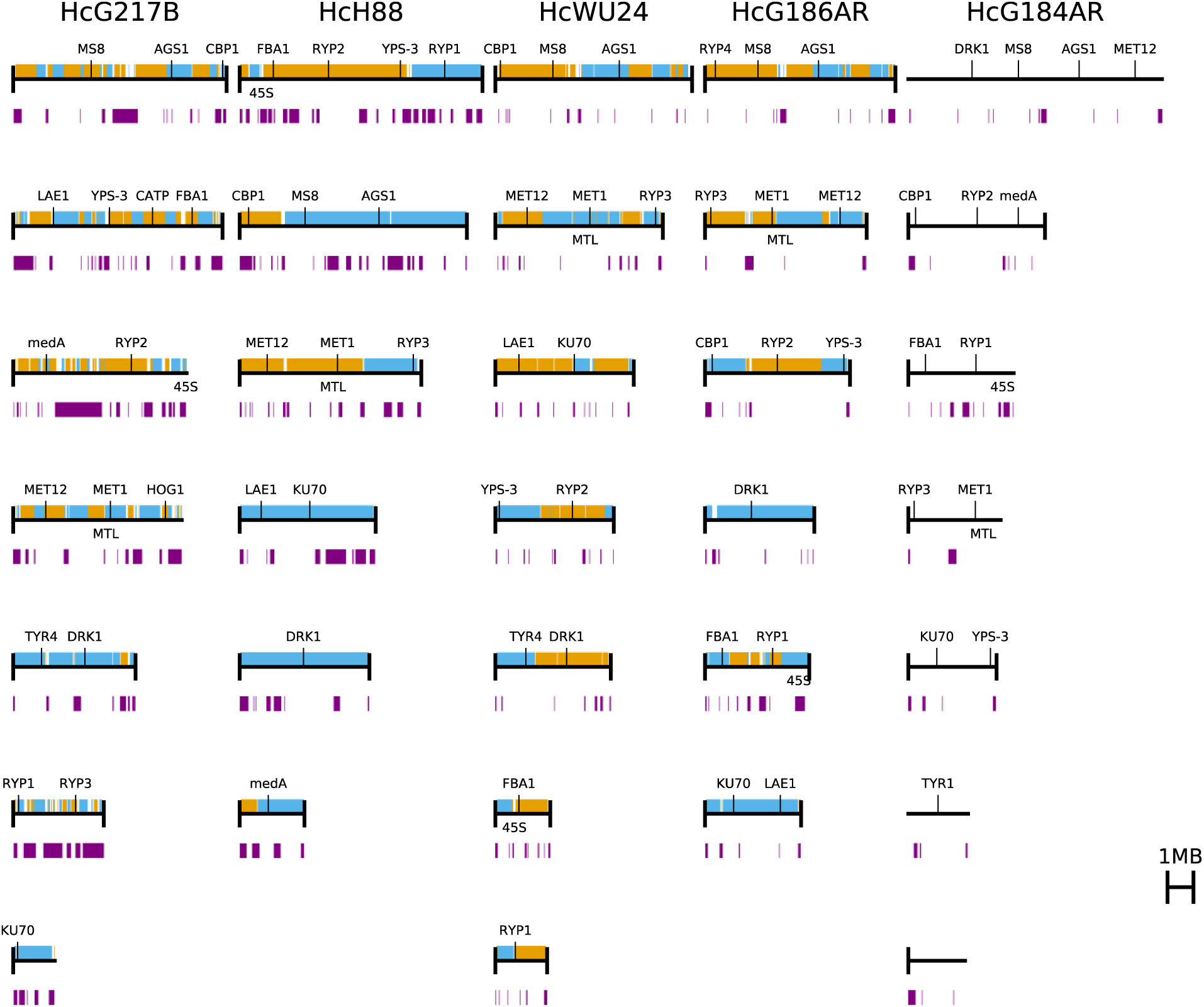
*Histoplasma* genomes are assembled at the chromosomal level. The newly assembled chromosomes of *Histoplasma* genomes are shown as black horizontal lines. Chromosomes or contigs are plotted largest to smallest. Telomeres are shown with black vertical lines at the end of each chromosome when present. Contigs from the previous assemblies are overlaid using alternating colors. Repeat regions (LTR transposons) are indicated with purple blocks below each chromosome. Genes of interest are displayed above each chromosome.

Next, we mapped existing gene annotations to the new assemblies using BLAT (17). For G217B, H88, G186AR, and G184AR strains, 88-92% of the RNA-Seq based annotations (18) -- using the G184AR annotations for both G184AR and G186AR -- transferred exactly and 96% transferred with no changes to the coding sequence (Table S2). For WU24, 77% of the previously available gene predictions transferred exactly and 81% transferred with no changes to the coding sequence.

### *Histoplasma* genomes have highly variable repetitive regions

The genomes of *Histoplasma* and *Blastomyces* have been observed to contain a substantial, but widely variable, amount of repetitive LTR transposon content (8, 19, 20). We annotated transposons using two complementary approaches: searching for broadly conserved features using LTRHarvest (21) and searching for protein coding regions homologous to the *gag* and *pol* open reading frames of a previously annotated *Histoplasma* LTR transposon using TBLASTN (22). The regions identified by the two methods had extensive overlap and were merged for further analysis. Results of our analysis revealed that the genomes of G217B and H88 are composed of 20% and 15% LTR transposon sequence respectively, compared to 4-5% for the other three genomes (Fig. 1). This difference in repeat content is sufficient to explain nearly all of the size differences among the genomes; after removing transposon sequence the genomes differ by at most 3 Mb (∼10% of the genome size). Visually, the transposon distribution appeared punctate rather than diffuse throughout the genome, which is reminiscent of the transposon distribution in *Blastomyces* (20). To examine this pattern further in *Histoplasma*, we joined transposons within 50 kb of each other. This analysis gave only 66-128 transposon-rich regions per genome, with the transposon-rich G217B having only 107 regions compared to 128 for the relatively transposon-poor WU24, confirming the punctate distribution of transposons within *Histoplasma* genomes.

### *Histoplasma* strains have highly syntenic chromosomes

Earlier studies showed that there is variability in chromosome size among *Histoplasma* strains (23, 24). Our results confirm these early findings and reveal that the repeat content is highly variable among the *Histoplasma* strains, which contributes to size differences. To investigate whether the gene order is conserved despite the chromosome size differences, we mapped the locations of orthologous genes among the *Histoplasma* genomes using our previous ortholog mappings for G217B, G186AR, and H88 (18) supplemented with INPARANOID (25) mappings to WU24, and assuming equivalent gene sets for the highly similar G186AR and G184AR strains. This ortholog-based synteny analysis gave results consistent with whole genome nucleotide alignments using NUCmer from MUMmer 3 (26). Our analyses showed that the genomes of H88, WU24, G217B, and G186AR are highly syntenic, with chromosomes 2, 3, and 5 of H88 essentially being conserved, with a few rearrangements, in the other strains (Fig. 2, S3-S7). Additionally, chromosome 4 of H88 is conserved in G186AR and is conserved but fused to different regions in G217B (with content orthologous to H88 chromosome 1) and WU24 (with content orthologous to H88 chromosome 6). Interestingly, G184AR is more dramatically rearranged relative to G186AR, which was isolated from the same Panama population and is highly identical at the nucleotide level (Fig. 2, S3-S7).

**Fig. 2.**
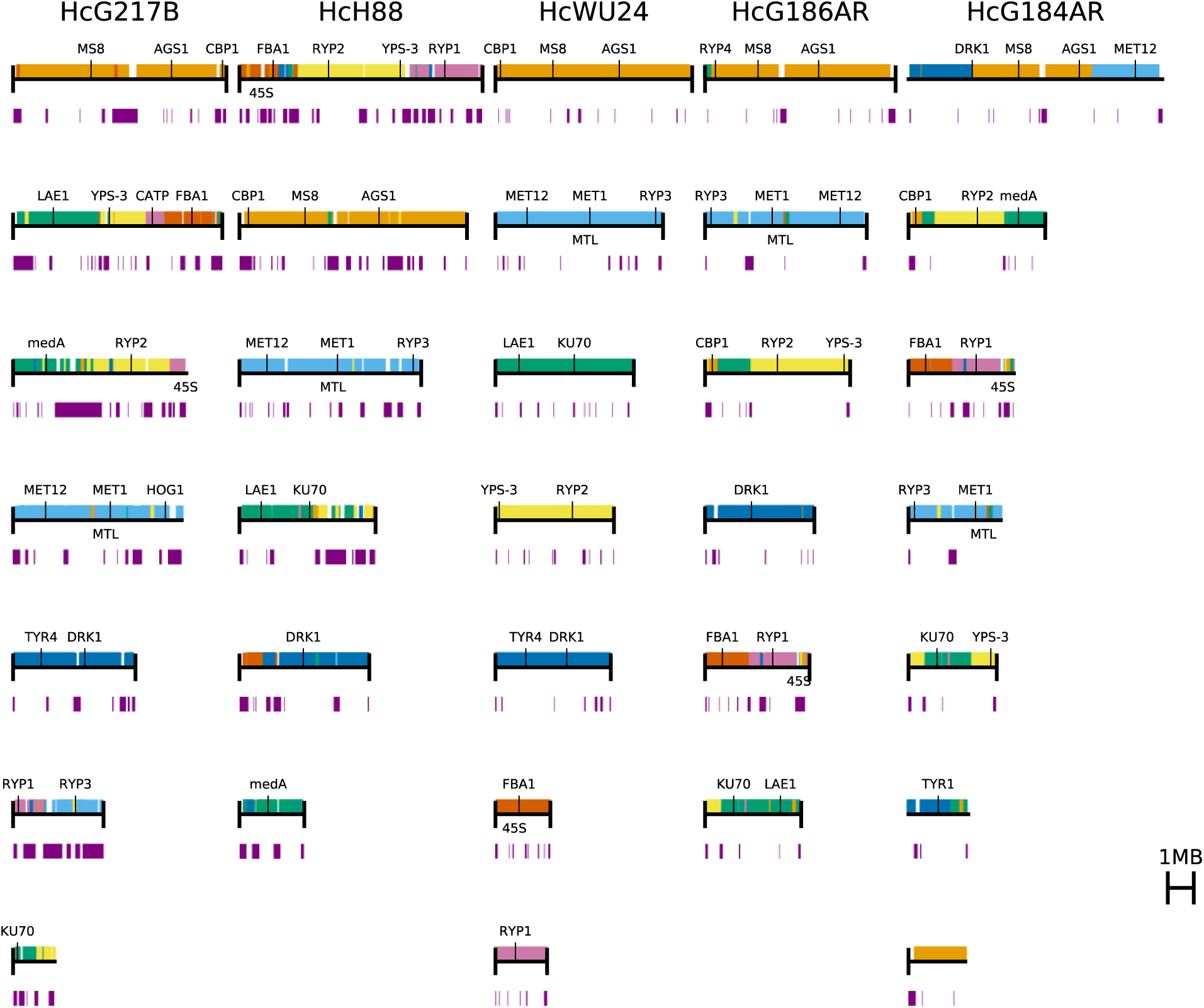
The *Histoplasma* genomes are highly syntenic. Chromosomes are sorted largest to smallest, and regions of synteny are colored according to the WU24 chromosomes in other strains. Purple bars below the chromosomes indicating the repetitive (transposon) regions.

Moreover, the locations of the transposon-rich regions are not conserved. Instead, the transposons appear to be inserted at idiosyncratic locations in each genome without large disruption of the conserved gene order (Fig. 2, S3-S8). Nevertheless, there does appear to be conservation of transposon-adjacent context for some genes. For example, the *CBP1* gene, encoding the most studied virulence factor of *Histoplasma* (27) has a conserved subtelomeric location that is flanked by transposons on the telomere side in WU24, G186AR, and G184AR, on both sides in G217B, and is embedded in a transposon-rich region in H88 (Fig. 2 and S9). The *CBP1* chromosome in WU24, G217B and H88 also contains *RYP4*, a regulator of morphology and dimorphism in *Histoplasma* (9).

An extensive synteny analysis between *Histoplasma* strains reveals conservation of gene order around many annotated genes (Fig. S9-S11). One of the most striking degrees of synteny can be observed in chromosome 3 of H88, which contains the mating type locus (28, 29) as well as most of the genes for sulfur assimilation, *SRE1*, a regulator of iron acquisition (30), and *RYP3*, a regulator of morphology and dimorphism in *Histoplasma* (*11*). This chromosome is syntenic along almost its entire length within the *Histoplasma* strains (Fig. S5 and S10). In addition, high conservation of the region containing *DRK1*, a key regulatory gene of dimorphism in *Blastomyces* and *Histoplasma* (*31*), is also remarkable among the *Histoplasma* strains and the closely related fungal pathogens *Blastomyces*, *Paracoccidioides* and *Coccidioides* (Fig. S11).

### Synteny analysis reveals duplication of a major regulator of fungal dimorphism

Our analysis also reveals the duplication of two ∼250 kb regions in G186AR relative to G184AR. G186AR and G184AR are the two Panamanian isolates and have a median identity of 99.95% for all large (>=100kb) alignable regions. The alignments were explored in more detail with BLASTN (Fig. 3B and C). Mapping of our G186AR Illumina reads to the G184AR assembly confirms ∼2x coverage in these regions (Fig. 3A), as does mapping of previously published G186AR Illumina reads (SRR6243650 (8)). Mapping of our G186AR ONT reads to the G184AR assembly likewise confirms the duplication; the boundaries of the duplicated regions have soft-clipping for about half of the mapped reads, as would be expected if these reads originate from the internal junction of the duplicated region (Fig. 3D and E).

**Fig. 3.**
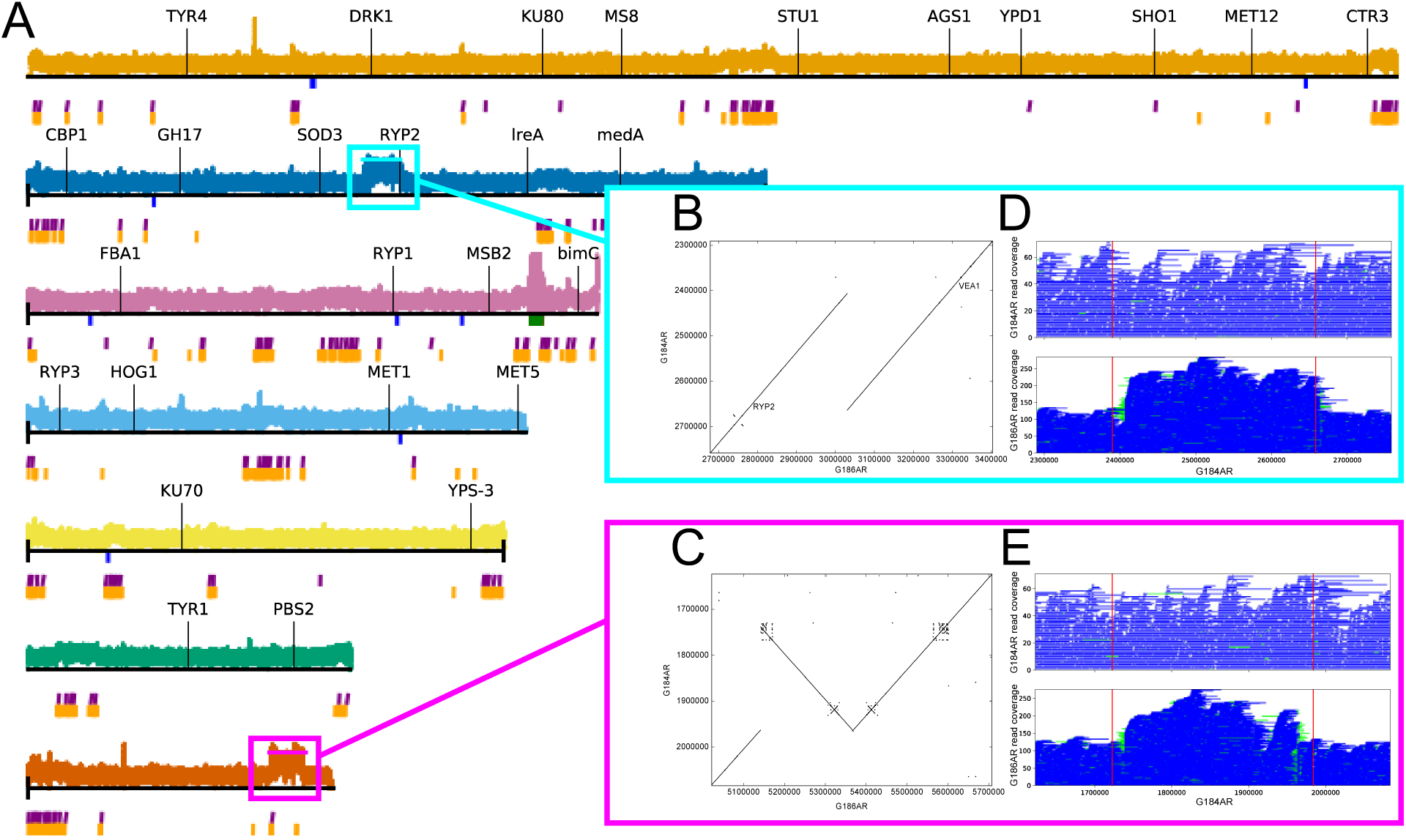
G186AR strain contains two duplicated regions. (A) Coverage for mapping G186AR Illumina reads onto the current G184AR assembly. (B,C) BLASTN-based dotplots for the duplicated regions. (D, E) Coverage for mapping G184AR and G186AR ONT reads onto the current G184AR assembly.

The first duplication (Fig. 3B) is a direct repeat from ∼16 kb to the left (3’) of *RYP2* to ∼8 kb to the left (5’) of *VEA1* (such that *RYP2* is duplicated but *VEA1*is not). *RYP2* and *VEA1* are both velvet transcription factors with roles in morphology in *Histoplasma* (11, 32). The duplicated region is not itself repetitive; it contains 110 gene annotations in G184AR (1 gene per 2.4 kb) and no LTR transposons (Fig. 3A).

The second duplication (Fig. 3C) is an inverted repeat. While mostly genic with ∼1 gene per 2.4 kb as for the first region, this second region does contain two sets of LTR transposons in G184AR. This duplicated region contains phospholipase B (*PLB1*) which was observed to have increased expression in G186AR yeast relative to G217B yeast (33). However, the same study also noted similar increased expression of phospholipase D and *OLE1*, which are not located in either duplicated region, so it is not clear if the copy number variation is an underlying cause of the expression difference. The yeast-enriched gene *TSA1* (GenBank AAK54753), a putative cysteine peroxidase, is also in the duplicated region.

### Transcripts embedded in transposon-rich regions are enriched in yeast phase

Fungal pathogens of plants contain transposon-rich genomic regions that are enriched for virulence factors with increased transcription in the host (34). In contrast, transposon-rich regions were not enriched for differential expression in *Blastomyces*, a close relative of *Histoplasma* (*20*). We performed similar analysis in *Histoplasma* using the previously published RNA-Seq datasets (18, 33) for four genomes, G217B, H88, G186AR and G184AR. We classified genes either as LTR (transposon), in (inside of the transposon-rich regions), near (within 50 kb of a transposon-rich region), or other (the remaining genes) based on their proximity to transposon -rich regions (Fig. 4, S12-S14 and Table S4). In all cases, the expression of transposon-adjacent genes is indistinguishable from transposon-distant genes (Fig. 4). In G217B and H88, genes annotated as transposons have increased expression in yeast compared to hyphae (Fig. 4). Due to the high sequence identity among transposons, we cannot distinguish whether this differential transcription occurs at some or all of the transposon loci. In G217B, the transposon-embedded genes likewise have increased expression in yeast compared to hyphae, significantly distinguishable from the transposon-adjacent and transposon-distant genes (p < 2.2e-16, Wilcoxon test). On closer inspection, this expression is distributed bimodally, with the transposon-embedded genes appearing to have expression ratios drawn either from the transposon or general distributions (Fig. 4). In H88, there is a less pronounced but still significant (p = 4.7e-15, Wilcoxon test) increase in yeast/hyphae expression for transposon-embedded genes. For the Panama strains, G186AR and G184AR, neither the transposons nor the transposon-adjacent genes have expression distributions distinguishable from the remaining genes (Fig. 4).

**Fig. 4.**
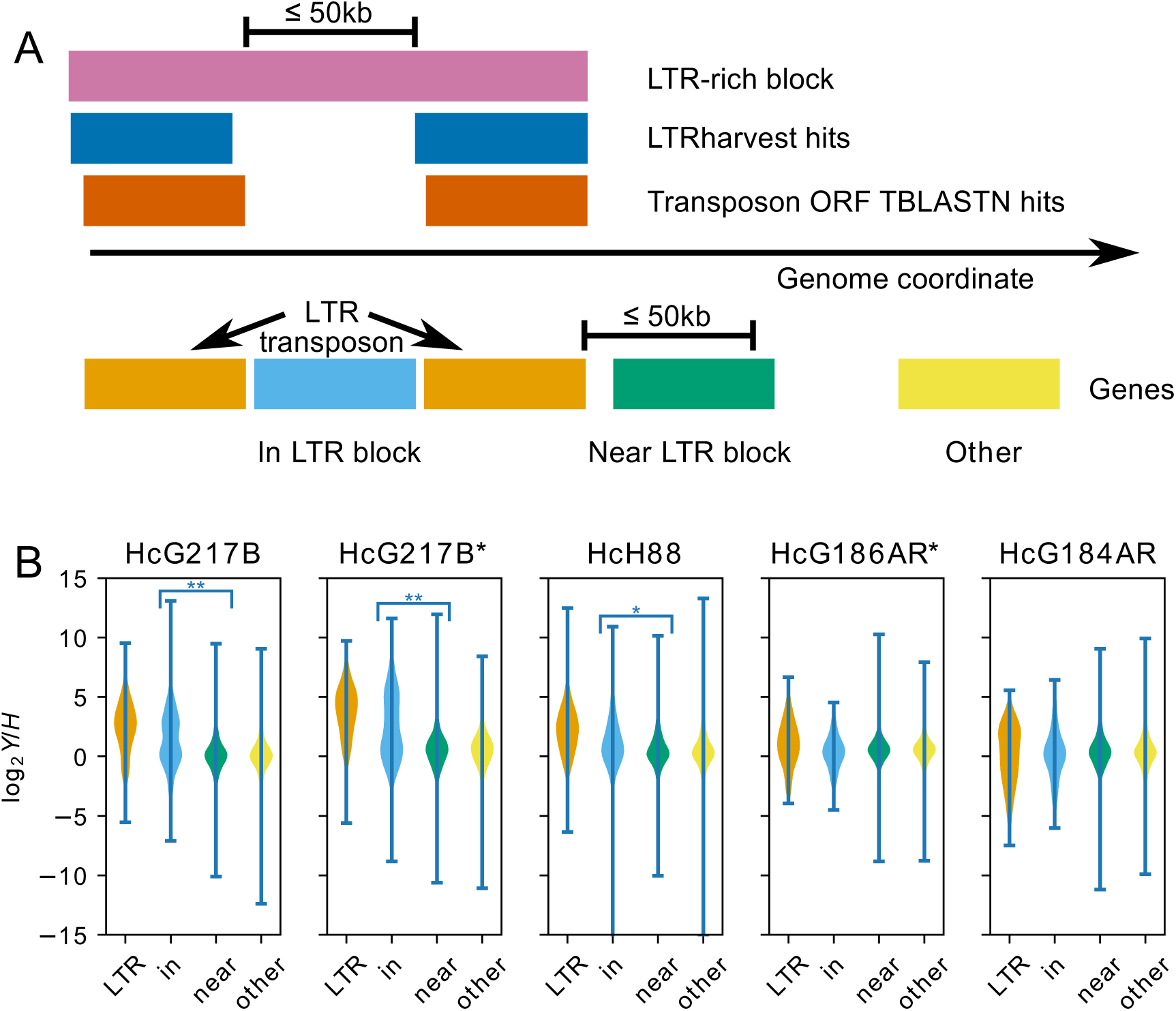
Transposon-embedded genes show yeast-phase enriched expression patterns. (A) Schematic of transcript classification by proximity to LTR transposon; *viz.*, “LTR” (orange): transcript overlaps transposon annotation, “in” (blue): transcript is between two transposon annotations within 50kb of each other, “near” (green): transcript is within 50kb of a transposon annotation, “other” (yellow): transcript does not fall into any of the previous categories. (B) Violin plots of yeast/hyphae expression ratios from (18) or (33)(*) taken from the Kallisto analysis of (12), colored according to the same scheme as A. Significant differences by Wilcoxon rank sum test are indicated: **: p < 2.2e-16, *: p = 4.7e- 15.

## Discussion

Advancement of sequencing technology, in particular the long reads produced by PacBio or ONT, has greatly improved our understanding of fungal genomes (13, 14, 37). Among over 1000 publicly available fungal genomes assemblies, there are currently 137 assemblies that have employed ONT sequencing. The most significant characteristic of these assemblies is that the N50 of Nanopore assemblies is about 1.4 Mb, whereas the N50 of all fungal assemblies is about 140 kb, suggesting at least a 10-fold improvement in the contiguity of the genome sequences by ONT sequencing (Fig. S15).

*Histoplasma* strains are important fungal pathogens with clinical significance. The previously available genomes of the most highly studied *Histoplasma* strains, G217B and G186AR, were highly fragmented. There are a number of analytical techniques (*e.g.*, evolutionary and taxonomic studies, transcriptional profiling, forward and reverse genetics, and mapping chemical DNA modifications and structural genomic changes) that would benefit from fully assembled genomes. Therefore, in this study, we have utilized ONT sequencing to fully assemble *Histoplasma* genomes.

With additional polishing using short reads from Illumina sequencing, we assembled genomes of five *Histoplasma* strains (WU24, G217B, G186AR, G184AR, and H88) at the chromosomal level. The sizes of the genomes vary from 31 Mb (G186AR and G184AR) to 40 Mb (G217B). The size differences among the strains were attributable to the amount of repeat content. After accounting for the repeat content, the non-repeat genome content (∼30 Mb) of *Histoplasma* is consistent with other Onygenales, such as *Coccidioides*, *Paracoccidioides*, *Blastomyces* and *Emmonsia* (20, 38–40).

Having near-complete chromosomes in hand, the first analysis we performed was investigating the synteny among *Histoplasma* strains, other close relatives (*Blastomyces dermatitidis*, *Paracoccidioides brasiliensis* and *Coccidioides immitis*) and a distant relative *Aspergillus nidulans*. Despite the variability in the repeat content among *Histoplasma* strains, the four strains WU24, G217B, G186AR and H88 are highly syntenic, whereas G184AR seems to be the most rearranged strain. This was an unexpected observation, as G186AR and G184AR are both Panamanian strains and are the most similar to each other at the nucleotide level. This indication that geographic location and nucleotide-level resemblance does not correlate with patterns of genomic rearrangement suggests that in some cases phylogenetic divergence may not be related to large-scale rearrangement. Additional synteny analyses around the previously studied genes (*viz.,* a virulence factor *CBP1*, sulfur assimilation genes, a regulator of iron acquisition *SRE1*, and a histidine kinase *DRK1*) showed a conservation of gene order throughout the *Histoplasma* strains. Moreover, a region around *DRK1* is highly syntenic throughout the Onygenales with substantial conserved gene order even in *Aspergillus* (Fig. S11). Chromosome-level assemblies revealed these cases of synteny in addition to larger-scale conserved regions that correspond to chromosomes 2, 3, 4, and 5 in H88.

Another important observation from the synteny analysis between *Histoplasma* strains was a duplication in G186AR of a region that contains the important developmental regulator *RYP2*. *RYP2* was previously identified as required for yeast phase in a screen in G217B (11) and, in complex with the velvet transcription factor *RYP3*, directly associates with promoters of yeast-enriched genes (9). The duplicated region also contains *HPD1*, a yeast-enriched gene (41) which has been shown to have a role in morphology in *P. brasiliensis* (42) and *Talaromyces marneffei* (43). Despite the duplication of these regulators in the G186AR strain, the yeast and hyphal transcriptional programs of G186AR (33) are grossly similar to those of G184AR (18), where the region appears to be in single copy. However, a detailed transcriptional analysis of the transition between yeast and hyphae has not been performed in these strains and could uncover kinetic differences that correspond to the duplication of developmental regulators.

One major advantage of having fully assembled genomes is to be able to visualize and analyze whole genome-level data (*e.g.,* transcriptional profiling) at the chromosomal level. Mapping of existing RNA-Seq datasets for G217B, G186AR, G184AR and H88 strains revealed that the transposon-adjacent genes are not distinguishable in their expression patterns from the transposon-distant genes. However, the transposon-embedded genes have significantly increased expression in yeast versus hyphae comparisons in two of the large, transposon-enriched *Histoplasma* genomes, viz; the G217B and H88 strains. While easily missed when analyzing the data relative to the previously available 261 contig G217B assembly, this large-scale pattern of transcriptional regulation is immediately apparent when plotted on the new near-chromosomal assembly. Further studies will reveal the importance of transposons-rich regions in *Histoplasma*. We predict that the new *Histoplasma* assemblies will likewise empower analysis of further genome-scale data sets going forward.

## Materials and Methods

### Sequencing of *Histoplasma* strains

Genomic DNA of *H. capsulatum* G217B, H88, G184AR, G186AR, and WU24 strains was harvested using phenol-chloroform extraction from yeast cultures grown to stationary phase in *Histoplasma* Macrophage Medium (44) at 37°C with 5% CO_2_ and shaking at 150 rpm. High molecular weight (HMW) gDNA was used to construct libraries for Illumina and ONT sequencing.

For G217B, G184AR, and H88, DNA libraries were generated using sequencing kit SQK-LSK108 and sequenced on the ONT MinION R9.4 flowcell (FLO-MIN106). For G217B, 207,770 reads were generated with an N50 of 24.7kb; for G184AR, 139,371 reads were generated with an N50 of 30.2kb; and for H88, 119,871 reads were generated with an N50 of 24.7kb. The raw fast5 files were basecalled using Guppy v3.2.2 (ONT) and the resulting fastq reads were utilized to build the assemblies following the pipelines most appropriate for each strain. Preliminary genome assemblies were created prior to the release of Guppy v3.2.2 and were built using the reads basecalled with Albacore v2.1.3 (ONT). However, all final assemblies only contain data from Guppy-basecalled reads. Additional libraries were generated with the TruSeq PCR-Free kit and sequenced on the Illumina MiSeq, generating paired-end 150 bp reads. Coverage for G217B, G184AR and H88 was 28X, 16X and 25X respectively.

For ONT sequencing of G186AR and WU24, libraries were constructed using the ligation sequencing kit (SQK-LSK109) and each library was loaded on a single flowcell (FLO-MIN106D) and run on a GridIon. Basecalling was performed with Guppy using MinKNOW 19.06.8. For WU24, a total of 578,143 reads were generated with a N50 of 29.2kb and estimated genome coverage of 202X. For G186AR, a total of 6,660,775 reads were generated with N50 of 4.8kb and estimated genome coverage of 456X. For Illumina sequencing of G186AR and WU24, 100 ng of genomic DNA was sheared to ∼250 bp using a Covaris LE instrument and prepared for sequencing as previously described (45); libraries were pooled and sequenced on a HiSeq2000 to generate paired 100 base reads, generating 150X coverage for WU24 and 152X coverage for G186AR.

### Genome Assemblies

Several methods of genome assembly were utilized to construct the final sequences (Fig. S2). The appropriate method for each genome was chosen by comparing the degree of fragmentation and read coverage. The MINIMAP/MINIASM/RACON (MMR) assembly protocol referenced below entails the following: ONT reads all-vs-all mapped using MINIMAP2 v2.13-r850 with parameter -x ava-ont (46). Overlapping regions assembled with MINIASM v0.3-r179 (47). ONT reads mapped back onto the overlap assembly using MINIMAP2 -x map-ont. RACON v1.3.1 (48) used to build a consensus assembly from this mapping and the ONT reads.

G217B ONT reads basecalled with Albacore were assembled following the MMR protocol, creating assembly *G217B_v1*. This assembly was polished using Illumina reads with 15 rounds of Pilon v1.22 (49). Assembly *G217B_v2* was created using the Guppy basecalled reads. These were first assembled using Flye v2.3.4 (50), then polished using Illumina reads with 16 rounds of Pilon. Alignment of *G217B_v1* and *G217B_v2* using NUCmer v3.1(26) indicated that one join could be made between two contigs in *G217B_v2*. Manually stitching this join resulted in assembly *G217B_v3*.

G184AR ONT reads basecalled with Albacore were assembled using wtdbg2 v2.3 and polished with Pilon using Illumina reads to create assembly *G184AR_v1*. Reads basecalled with Guppy were assembled following the MMR protocol, and polished with Pilon using Illumina reads to create assembly *G184AR_v2*. *G184AR_v1* was used to guide contig joins on *G184AR_v2*, producing assembly *G184AR_v3*.

H88 ONT reads basecalled with Albacore were assembled using Flye v2.3.4 to construct the genomic DNA sequence *H88_v1*. The mitochondrial DNA sequence *H88_v2* was built from the same reads assembled using wtdbg2 v2.3, then polished with 15 rounds of Pilon using Illumina reads. These two assemblies were combined to form *H88_v3*. Assembly *H88_v4* was created with the Guppy basecalled reads following the MMR protocol, then polished with four rounds of Pilon using the Illumina reads. *H88_v3* and *H88_v4* were aligned with NUCmer v3.1, which implied one join in *H88_v4*. This join was manually assembled to create *H88_v5*.

Contigs from *G217B_v3*, *G184AR_v3*, and *H88_v5* were then extended through the telomeres. This was accomplished by using minimap2 to align the ONT reads to the contigs. Reads containing telomeric repeats which extended past the ends of the contigs were extracted, and wtdbg2 (51) was used to assemble the telomeric ends of each chromosome.

G186AR and WU24 ONT reads were assembled into *G186AR_v1* and *WU24_v1* using Canu (version v1.6 with parameters genomeSize=30000000 stopOnReadQuality=false correctedErrorRate=0.075) (52). The assemblies were inspected and refined using alignments of contig ends, spanning reads, and to assemblies *G186AR_v2* and *WU24_v2* generated using Flye (version 2.7b-b1526 with parameter--genome-size=30000000). These alignments were used to identify and fix mis-assemblies (including building out the second copies of two large collapsed duplications in G186AR), extend contig ends to telomeric repeats, and make joins between contig ends. The mitochondrial contig in WU24 appears complete based on end overlap, which was trimmed. This refinement produced assemblies *G186AR_v3* and *WU24_v3*.

To produce *G217B_final*, *G184AR_final*, *H88_final*, *G186AR_final*, and *WU24_final*, all five preliminary assemblies were polished first with Medaka (version 0.8.1, using only the Nanopore reads) and then the Medaka-polished contigs were polished using Illumina reads aligned with BWA mem and Pilon (version 1.22 and 1.23), with default settings for both Medaka and Pilon. *G217B_final*, *G184AR_final*, *H88_final*, *G186AR_final*, and *WU24_final* underwent 6, 5, 6, 3, and 3 rounds of Pilon respectively.

### Assembly Quality Control

Mitochondrial DNA contigs in all assemblies were determined to be complete based on end overlap indicating circularity. Genome assembly quality and completeness was assessed using BUSCO version 4.0.4 (53) with dataset eurotiomycetes_odb10.

### Gene annotation

Transcript sequences for G217B, H88, and G184AR were taken from Gilmore et al. (18). The G184AR transcripts were used for G186AR as well. Transcript sequences for WU24 were downloaded from the Broad Institute on 6/15/2011 and are available at https://histo.ucsf.edu/downloads/histoplasma_capsulatum_nam1_1_transcripts.fasta

Transcripts were mapped to the new assemblies using BLAT version 35 invoked as blat -q=dna -t=dna -fine -noTrimA -maxIntron=10000 -minIdentity=98 retaining the top scoring hit for each transcript. Genes with CDS corrupted by the BLAT transfer were repaired, where possible, based on TBLASTN of the original protein sequence to the new genome assembly prior to submission to GenBank.

### Transposon annotation

Transposons were identified by mapping the translated open reading frames of a representative full length MAGGY transposon to each genome assembly by TBLASTN from NCBI BLAST+ 2.6.0 with default parameters. Transposons were additionally identified by LTRharvest from GenomeTools 1.5.9 with default parameters. For the purpose of classifying genes by proximity to transposons, transposon-rich regions were defined as contiguous bases within 50kb of a TBLASTN or LTRharvest identified transposon.

### Synteny analysis

Genes orthologous among G217B, H88, and G184AR were taken from Gilmore et al. Orthologs between G184AR and G186AR were assigned based on top BLAT hits to the same query sequence, as described above in “Gene annotation”. Orthologs between WU24 and the remaining genomes were assigned by INPARANOID 1.35.

Global synteny patterns were explored by coloring genes according to their chromosome in a reference genome (as in Fig. 2 and Fig. S4-S7) and by ortholog-based dotplots (as in Fig. S3). This method gave comparable results to nucleotide based dotplots generated by NUCmer from MUMmer 3.23.

Detailed synteny plots for a given ordered set of genomes and a given query region on the first genome (as in fig. S8-S11) were generated as follows: we first identified all genes in the query region with an ortholog in all of the remaining genomes. Then, for each genome, we identified all regions with at least 5 such orthologs separated by no more than 500kb. These regions were then plotted in alternating colors with lines connecting orthologous genes between adjacent genomes. The plotted regions were oriented to minimize crossovers between the ortholog-connecting lines.

### Analysis of G186AR duplications

Two large (∼250 kb) duplications in G186AR were initially observed as mapping artifacts in MINIMAP2 mapping of ONT reads (as MINIMAP2 generates incomplete mappings for reads with multiple high identity matches in the target genome). These duplications were also evident in NUCmer dotplots of G186AR vs. G184AR and were explored in detail with BLASTN. ∼2x coverage of the unduplicated G184AR regions by G186AR reads were confirmed using BWA MEM 0.7.15 for Illumina reads and MINIMAP2 (git commit d90583b83cd81a) for ONT reads.

### Fungal phylogeny analysis

The Pfam Gcd10p domain of Gcd10p was identified in each gene set using HMMSEARCH and aligned with HMMALIGN from HMMER3. FASTTREE2 was used to estimate a phylogeny from the resulting protein multiple alignment.

### Fungal genome query

Two searches were executed on October 19, 2020 in order to determine the contig N50 for all fungal genomes compared to fungal genomes sequenced using Oxford Nanopore technology. Fungal genome assemblies were queried by searching NCBI genome records with the search term *fungi[Organism]*. 1,395 related assembly records were extracted from this search. ONT-based fungal genome assemblies were queried by searching NCBI assembly records with the search term *fungi[Organism] AND nanopore[Sequencing Technology]*. The output of this search was 137 total assembly records. The assembly record XML summaries were downloaded from NCBI and the contig N50 values were extracted from them. The median contig N50 for the 1,395 fungal assemblies was 140,278 bp. The median contig N50 for the 137 Nanopore fungal assemblies was 1,400,166 bp.

### Data availability

Sequencing reads and annotated genome assemblies were submitted to GenBank under BioProjects PRJNA682643, PRJNA682644, PRJNA682645, PRJNA682647, and PRJNA682648.

## Acknowledgements

We thank the J. Craig Venter Institute (JCVI) Sequencing Core for generating Illumina and ONT reads, the Broad Institute Genomics Platform for generating Illumina sequences, and Ross Ackerman and the Broad Microbial ‘Omics Core for generating ONT reads for this study. We are grateful to Chris Dupont and Drishti Kaul for their help with bioinformatics at JCVI.

---

**Table S1.** Statistics of previous and current Histoplasma genome assemblies. Completeness of the genome assemblies was assessed using BUSCO version 4.0.4, with dataset eurotiomycetes_odb10 (53). Statistics and percentage of complete BUSCO groups are shown for previously published genomes and the assemblies reported in this study.

| Strain | Source | Method | Length | Contigs | Ctg N50 | Ctg L50 | Scaffolds | Sc N50 | Sc L50 | BUSCO |
| --- | --- | --- | --- | --- | --- | --- | --- | --- | --- | --- |
| WU24 | BROAD | Sanger | 33,030,326 | 2,873 | 27,346 | 233 | 280 | 2,885,091 | 4 | 87.4% |
| WU24 | This study | Oxford Nanopore | 32,531,515 | 8 | 5,328,076 | 3 | -- | -- | -- | 98.8% |
| G217B | WUSTL | Sanger | 39,270,953 | 261 | 634,562 | 21 | -- | -- | -- | 98.6% |
| G217B | This study | Oxford Nanopore | 39,447,273 | 12 | 6,680,169 | 3 | -- | -- | -- | 98.6% |
| G186AR | BROAD | Sanger | 30,483,324 | 359 | 191,715 | 48 | 88 | 1,769,553 | 6 | 97.4% |
| G186AR | This study | Oxford Nanopore | 31,111,494 | 7 | 5,585,295 | 3 | -- | -- | -- | 98.8% |
| G184AR | This study | Oxford Nanopore | 30,991,520 | 12 | 4,063,218 | 3 | -- | -- | -- | 98.8% |
| H88 | BROAD | Sanger | 37,943,219 | 464 | 168,190 | 67 | 17 | 5,001,999 | 4 | 97.7% |
| H88 | This study | Oxford Nanopore | 37,996,987 | 7 | 7,000,705 | 3 | -- | -- | -- | 98.9% |
| H143 | BROAD | Sanger | 38,961,716 | 4,267 | 15,865 | 663 | 49 | 1,740,271 | 7 | 85.0% |

**Table S2.** Statistics of annotation transfer for the new Histoplasma genome assemblies. Number of transcripts that either failed to map by BLAT to the new assembly (Not mapped), mapped but with a change in coding sequence length (CDS len), with a change in coding sequence (CDS seq), with a change in UTR length (Exon length), a change in UTR sequence (Exon seq), or with no changes to CDS or UTR (Match). Each transcript is recorded in the leftmost applicable column, such that the sum over each row gives the total previously annotated transcripts for that genome.

| Genome | Not mapped | CDS len | CDS seq | Exon length | Exon seq | Match |
| --- | --- | --- | --- | --- | --- | --- |
| WU24 | 54 (1%) | 385 (4%) | 1397 (15%) | 187 (2%) | 145 (2%) | 7080 (77%) |
| G217B | 31 (0%) | 12 (0%) | 443 (4%) | 843 (7%) | 104 (1%) | 10880 (88%) |
| G186AR | 11 (0%) | 5 (0%) | 348 (3%) | 529 (4%) | 123 (1%) | 11647 (92%) |
| G184AR | 31 (0%) | 10 (0%) | 527 (4%) | 819 (6%) | 117 (1%) | 11159 (88%) |
| H88 | 15 (0%) | 15 (0%) | 457 (4%) | 546 (4%) | 80 (1%) | 11062 (91%) |

**Table S3.** Analysis of repeat regions in *Histoplasma* genomes. Transposon statistics for the ONT genome assemblies are shown. Total = total base pairs, Repeat = number of base pairs spanned by union of TBLASTN and LTRHarvest annotations, Annealed = number of base pairs spanned by LTR-rich blocks (*c.f.* Fig. 3A), T-R = Total - Repeat, T-A = Total - Annealed, R% = Repeat/Total, A% = Annealed/Total, RL = number of repeat annotations (after union), AL = number of LTR-rich blocks, tgenes = number of transposon genes, rgenes = number of transposon-embedded genes, agenes = number of transposon-adjacent genes, %r = rgenes/(total genes), %a = agenes/(total genes).

| Genome | Total | Repeat | Annealed | T-R | T-A | R% | A% | RL | AL | tgenes | rgenes | agenes | %r | %a |
| --- | --- | --- | --- | --- | --- | --- | --- | --- | --- | --- | --- | --- | --- | --- |
| HcG217B | 39447273 | 7911417 | 13807944 | 31535856 | 25639329 | 0.2 | 0.35 | 1991 | 95 | 12282 | 371 | 1133 | 0.03 | 0.09 |
| HcH88 | 37996987 | 5892053 | 11804992 | 32104934 | 26191995 | 0.16 | 0.31 | 1609 | 87 | 12160 | 165 | 775 | 0.01 | 0.06 |
| HcWU24 | 32531515 | 1148983 | 3213981 | 31382532 | 29317534 | 0.04 | 0.1 | 387 | 128 | 9194 | 45 | 403 | 0 | 0.04 |
| HcG186AR | 31111494 | 1717918 | 3400858 | 29393576 | 27710636 | 0.06 | 0.11 | 493 | 70 | 12652 | 202 | 484 | 0.02 | 0.04 |
| HcG184AR | 30991520 | 1710517 | 3311154 | 29281003 | 27680366 | 0.06 | 0.11 | 480 | 69 | 12632 | 202 | 449 | 0.02 | 0.04 |

**Table S4. Histoplasma orthogroups.** A table of orthogroups with one row per orthogroup from the mappings used in the synteny analysis. Columns are: (1-2) short gene names and descriptions taken from Gilmore et al. with updates from Rodriguez et al., (3-7) systematic gene name (from the previous Broad annotation for WU24 or the Gilmore et al paired-end assembly for the other genomes), (8-12) repeat classification, as in Fig. 4A., (13-19) mappings to previous Broad and WUSTL predicted gene sets (_pred) or the Edwards et al transcriptomes assembled from unstranded RnaSeq (_unstranded), and (20-24) locus tags for genes submitted to GenBank. Previous gene sets were mapped to the new assemblies by BLAT and then to the current gene sets based on same-strand overlap. Where two previous genes mapped to the same location due to redundant sequence, the lexically first gene was chosen. One-to-many overlaps between different gene sets were resolved by choosing the gene pair with the greatest in-frame coding sequence overlap.

**Fig. S1.**
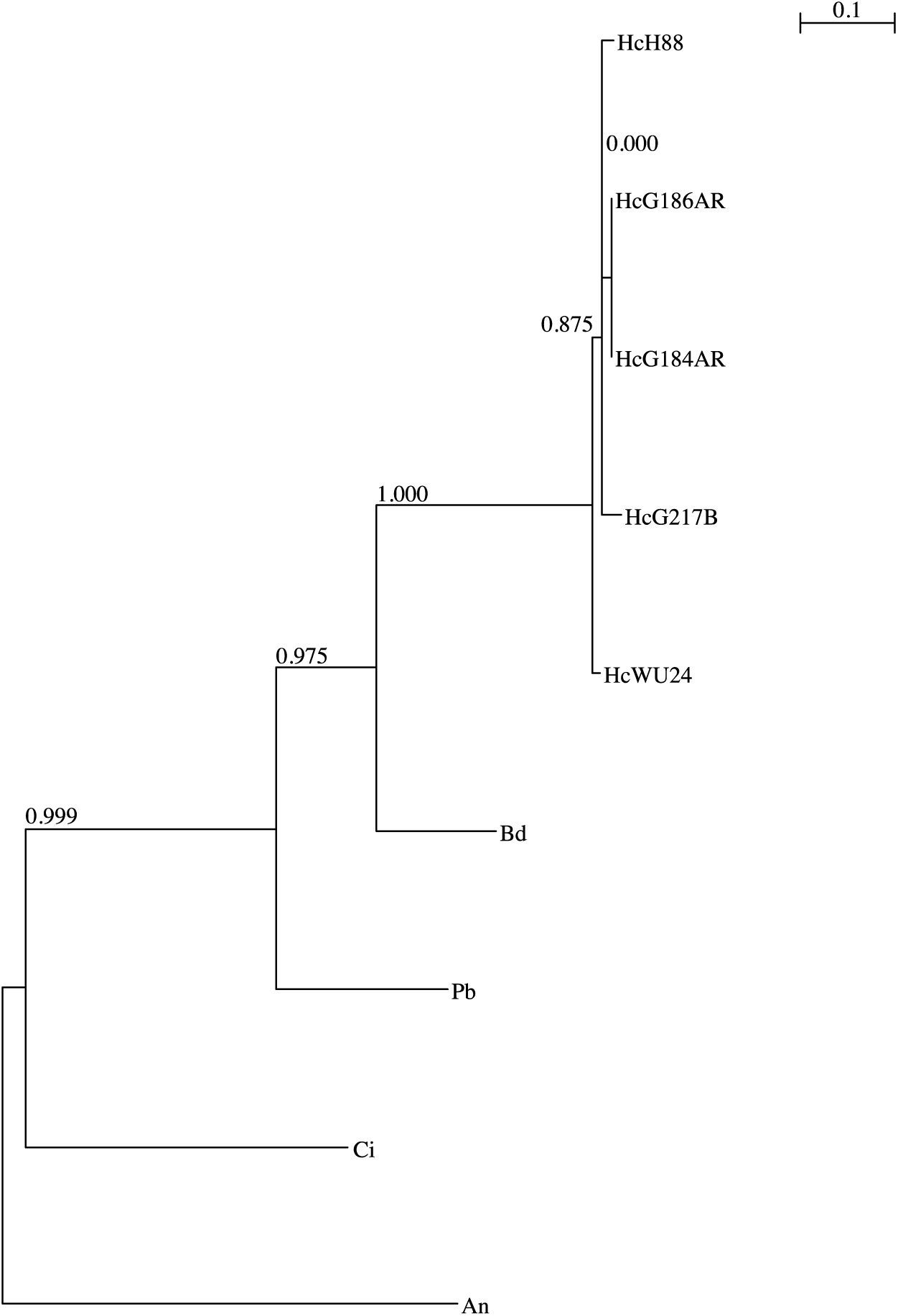
Phylogeny of Gcd10p from *Histoplasma* and related species. The Pfam Gcd10p domain of Gcd10p was identified in each gene set from five *Histoplasma* strains (HcH88, HcG186AR, Hc184AR, HcG217B, HcWU24), *Blastomyces dermatitidis* (Bd), *Paracoccidioides brasiliensis* (Pb), *Coccidioides immitis* (Ci) and *Aspergillus nidulans* (An) using HMMSEARCH and aligned with HMMALIGN from HMMER3. FASTTREE2 was used to estimate a phylogeny from the resulting protein multiple alignment. Numbers indicate FASTTREE2 bootstrap estimates.

**Fig. S2.**
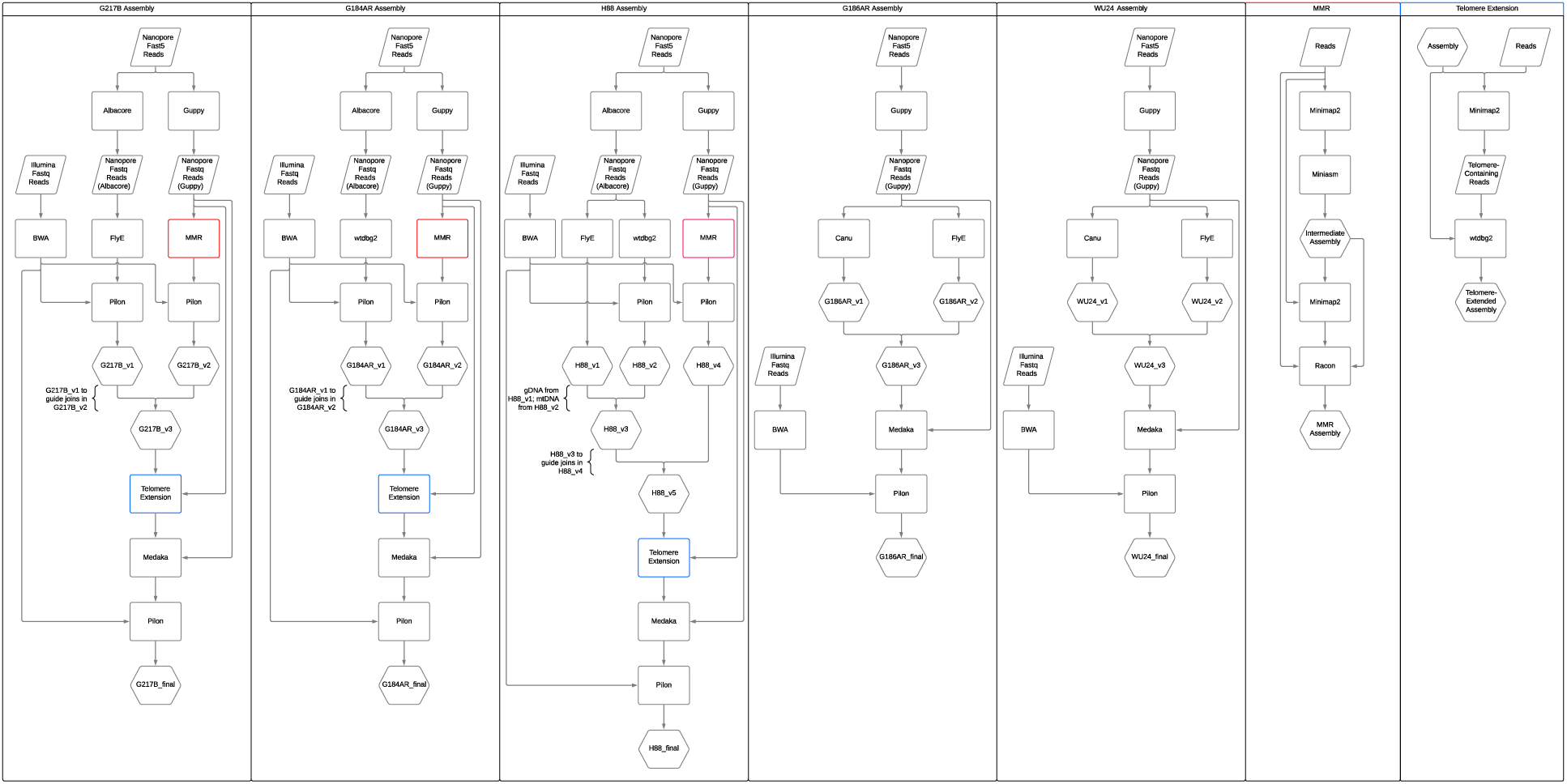
Fungal genome assembly pipeline. Assembly methods for the five genomes in this study are shown. MMR and Telmore Extension methods are expanded in separate columns.

**Fig. S3.**
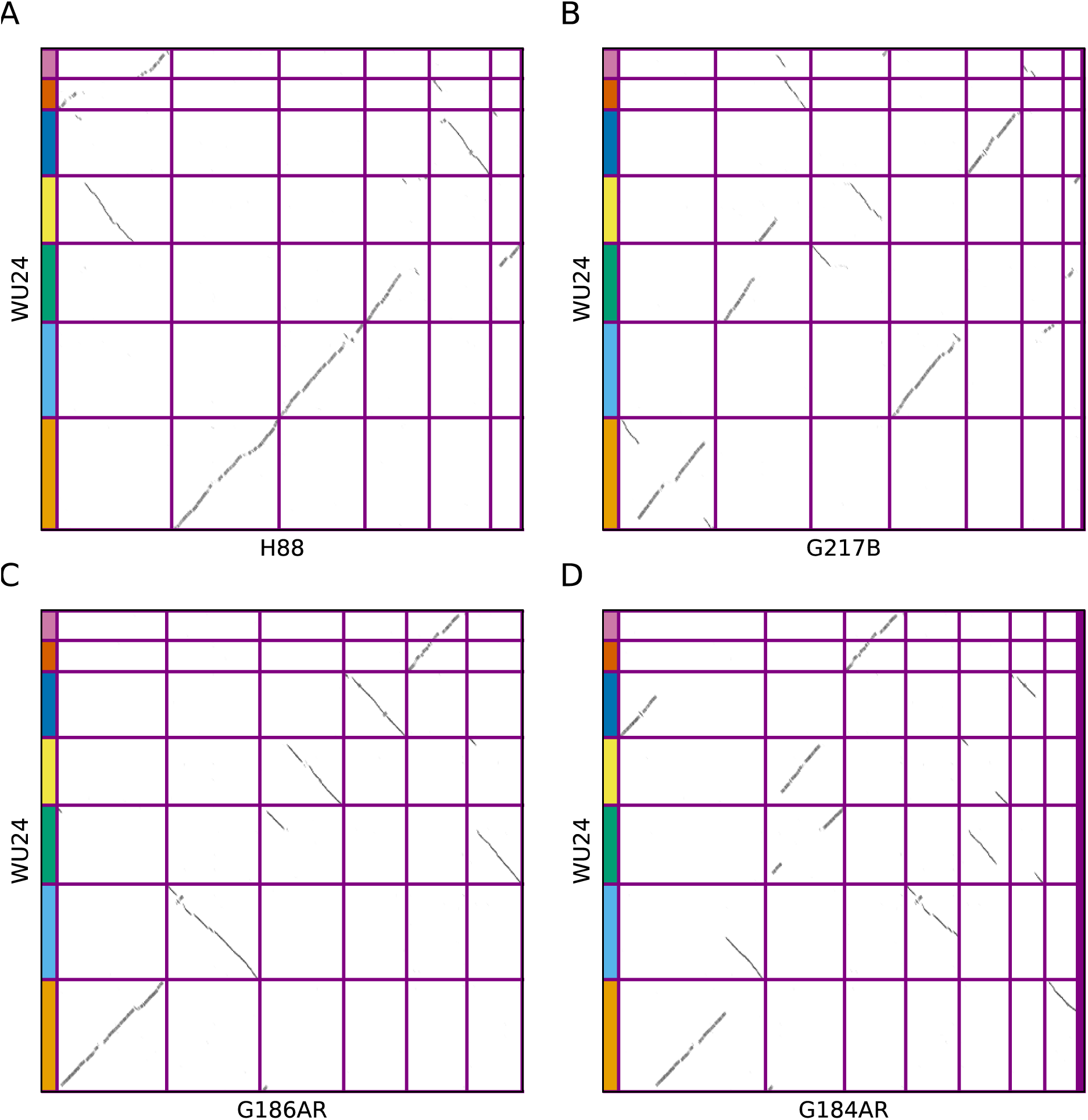
The *Histoplasma* genomes are highly syntenic. Dotplots showing genomic coordinates of orthologous genes between WU24 (y axis) and H88 (a), G217B (b), G186AR (c), and G184AR (d). Chromosomes are sorted largest to smallest, with purple lines indicating chromosome/contig boundaries. Genes are colored according to WU24 chromosome, as in Fig. 2.

**Fig. S4.**
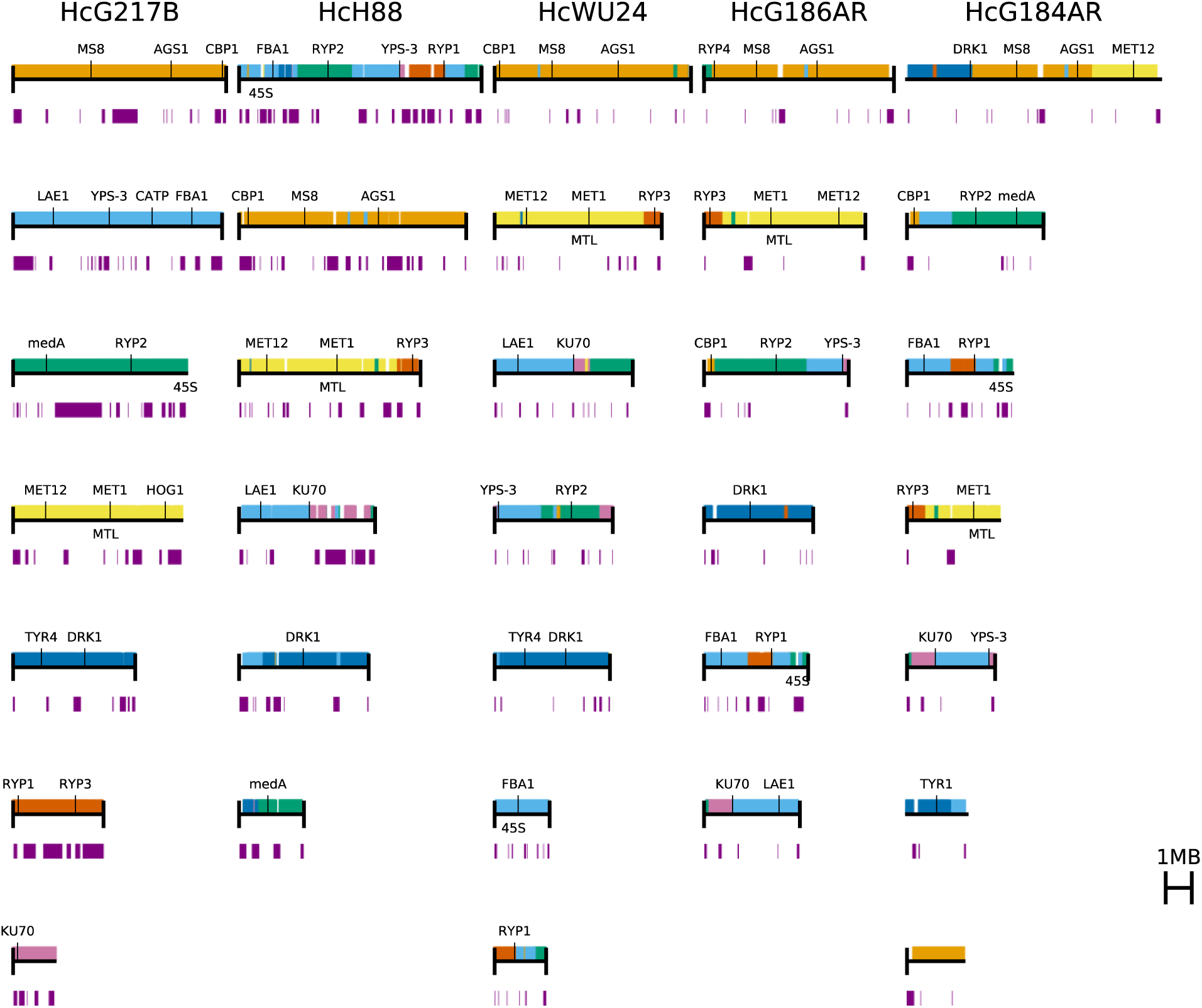
Synteny of the *Histoplasma* genomes relative to G217B. Chromosomes are sorted largest to smallest, and genes are colored according to the G217B chromosomes in other strains. Purple bars below the chromosomes indicating the repetitive regions.

**Fig. S5.**
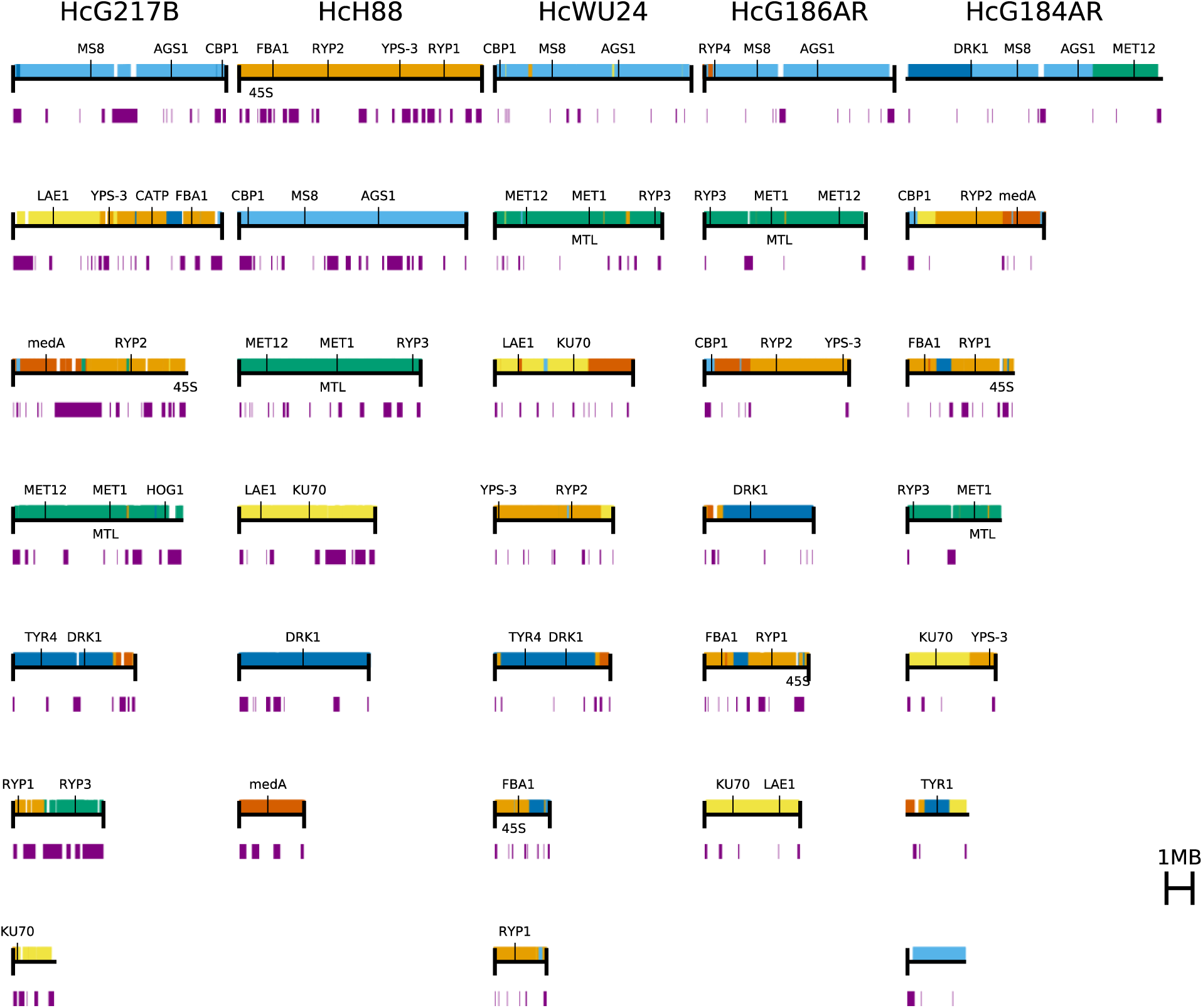
Synteny of the *Histoplasma* genomes relative to H88. Chromosomes are sorted largest to smallest, and genes are colored according to the H88 chromosomes in other strains. Purple bars below the chromosomes indicating the repetitive regions.

**Fig. S6.**
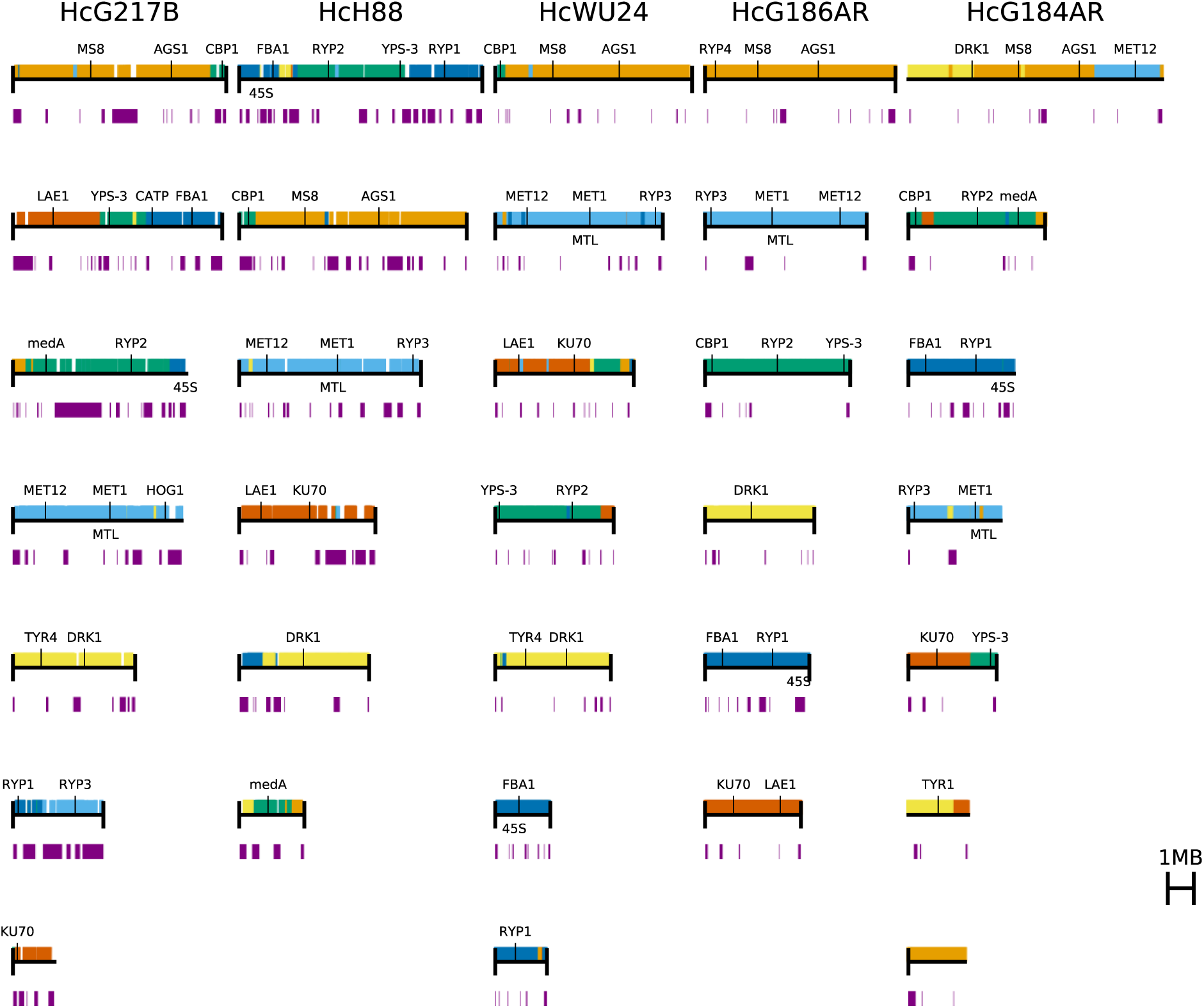
Synteny of the *Histoplasma* genomes relative to G186AR. Chromosomes are sorted largest to smallest, and genes are colored according to the G186AR chromosomes in other strains. Purple bars below the chromosomes indicating the repetitive regions.

**Fig. S7.**
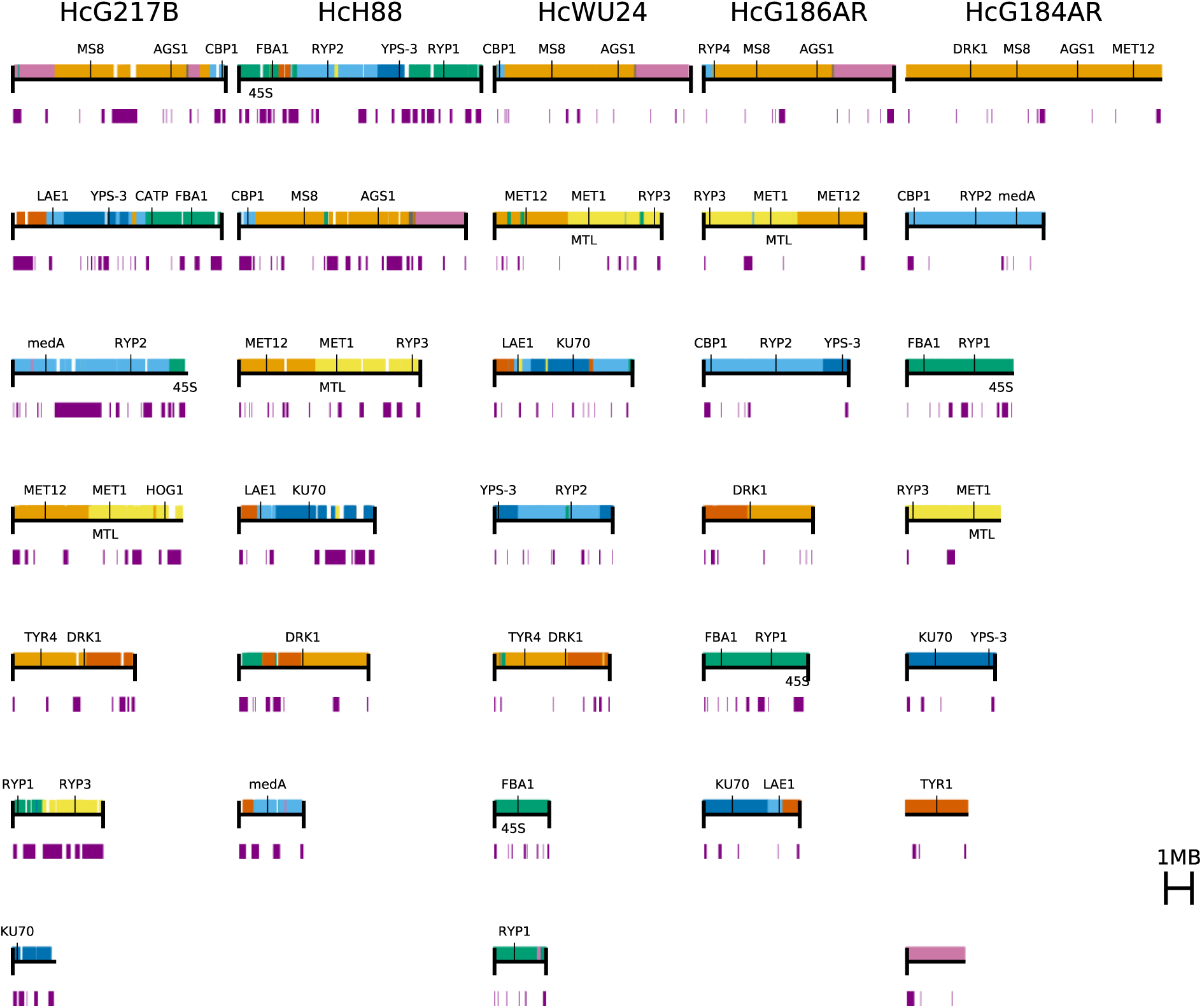
Synteny of the *Histoplasma* genomes relative to G184AR. Chromosomes are sorted largest to smallest, and genes are colored according to the G184AR chromosomes in other strains. Purple bars below the chromosomes indicating the repetitive regions.

**Fig. S8.**
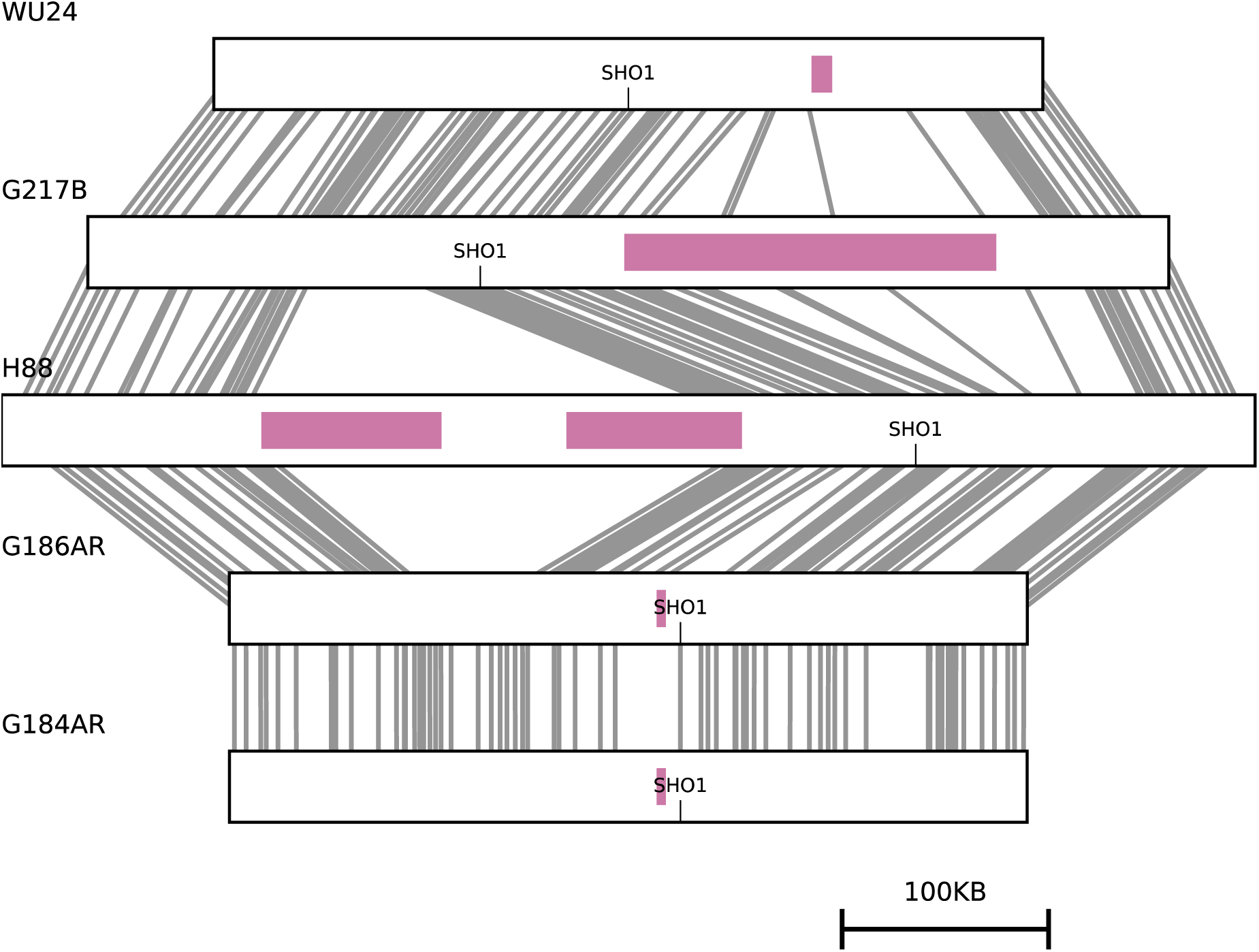
*SHO1* is in a syntenically conserved subtelomeric region. *SHO1* region of WU24 aligned to syntenic regions of selected genomes. Lines connect complete orthogroups across all genomes. Syntenic regions are regions with at least five genes orthologous to the WU24 region with no more than 500 kb between orthologous genes. Repetitive regions are shown as pink boxes inside of the syntenic regions.

**Fig. S9.**
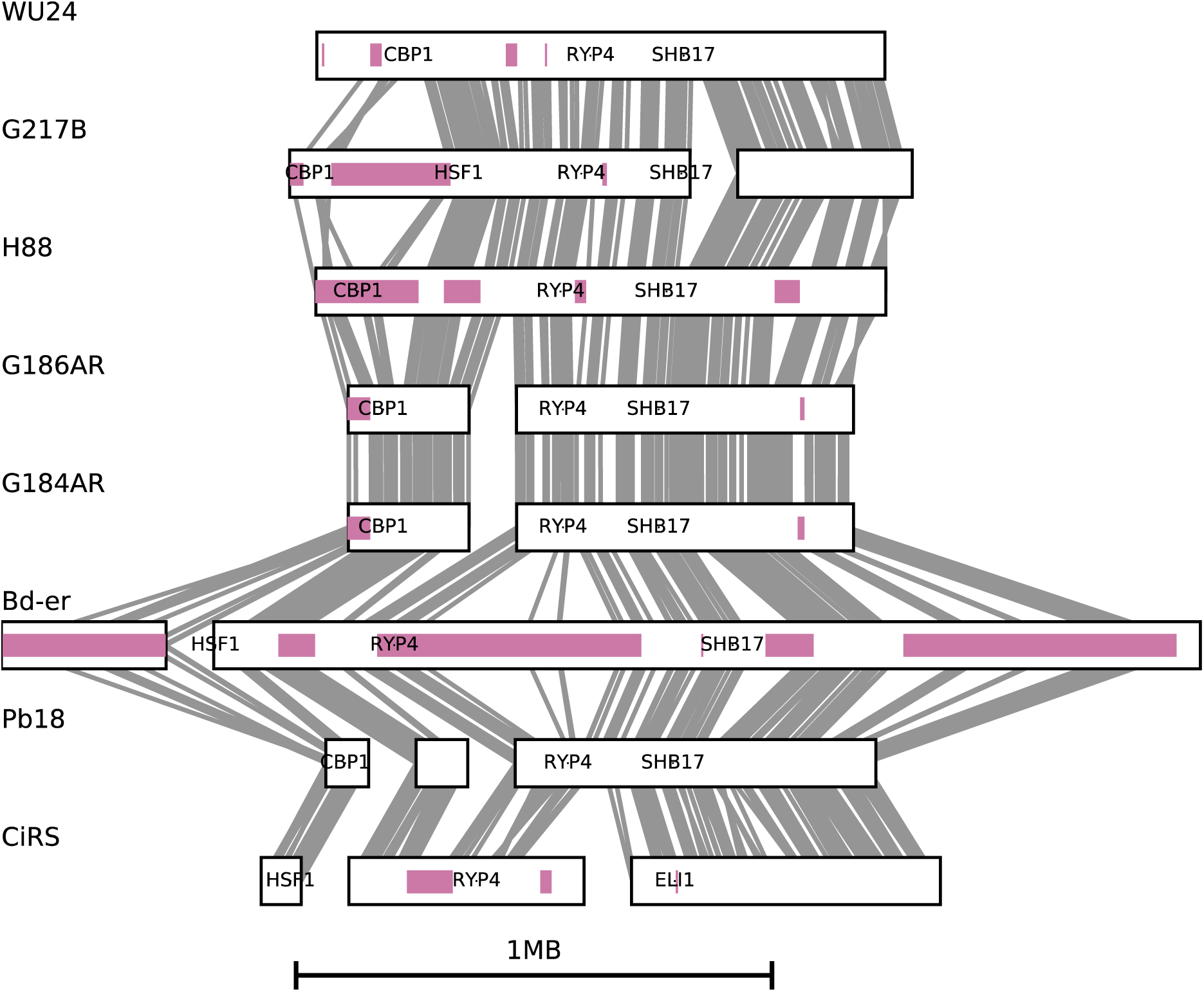
*CBP1* is in a syntenically conserved subtelomeric region. 1 Mb *CBP1*- containing subtelomeric region of WU24 aligned to syntenic regions of selected genomes. Lines connect complete orthogroups across all genomes. Syntenic regions are regions with at least five genes orthologous to the WU24 region with no more than 500 kb between orthologous genes. For genomes with broken synteny (syntenic regions on distinct chromosomes or separated by more than 500 kb) the separate syntenic regions are plotted next to each other. Repetitive regions are shown as pink boxes inside of the syntenic regions.

**Fig. S10.**
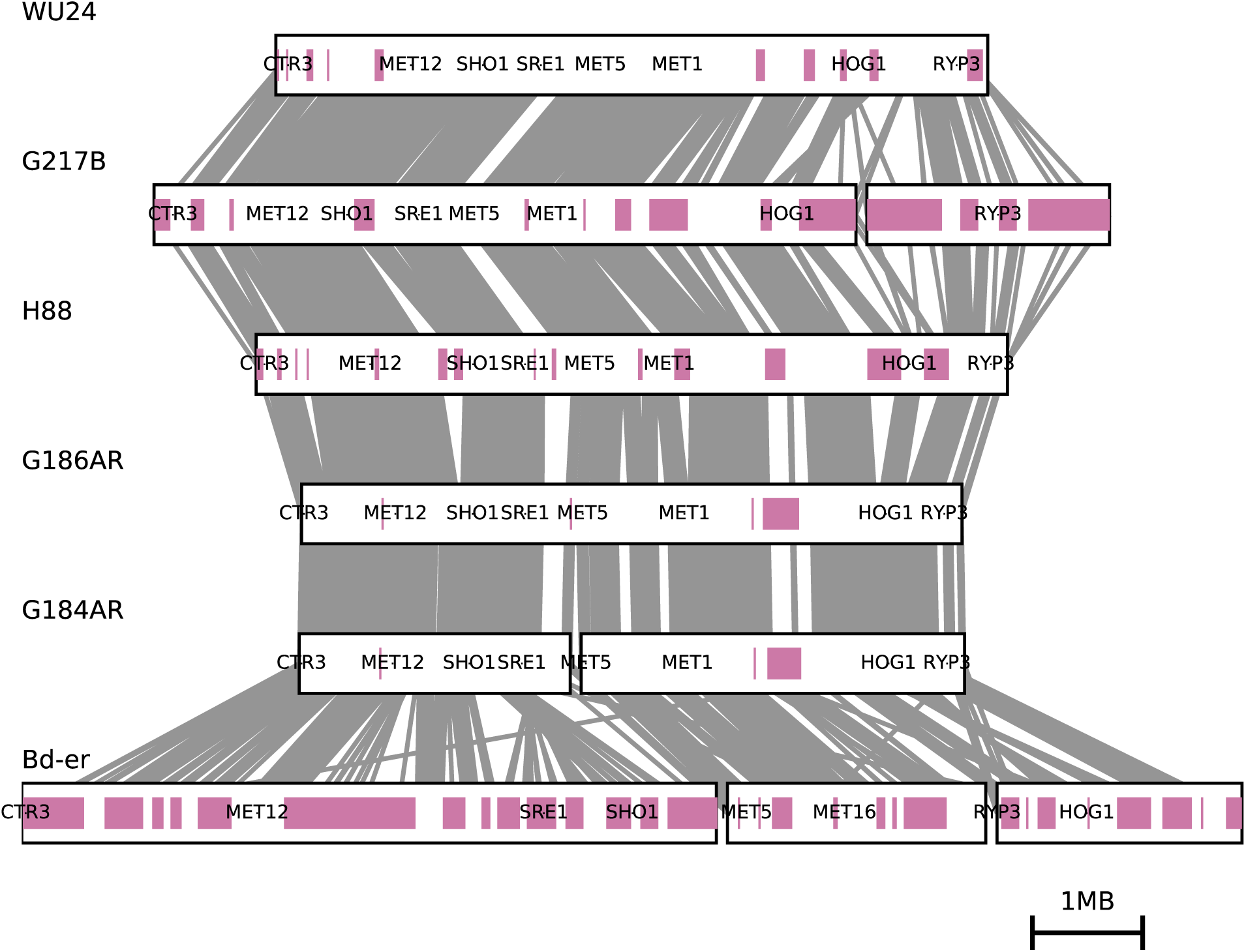
Sulfur assimilation genes are syntenic in *Histoplasma*. Chromosome 2 of WU24 aligned to syntenic regions of selected genomes, rendered as in Fig. S9.

**Fig. S11.**
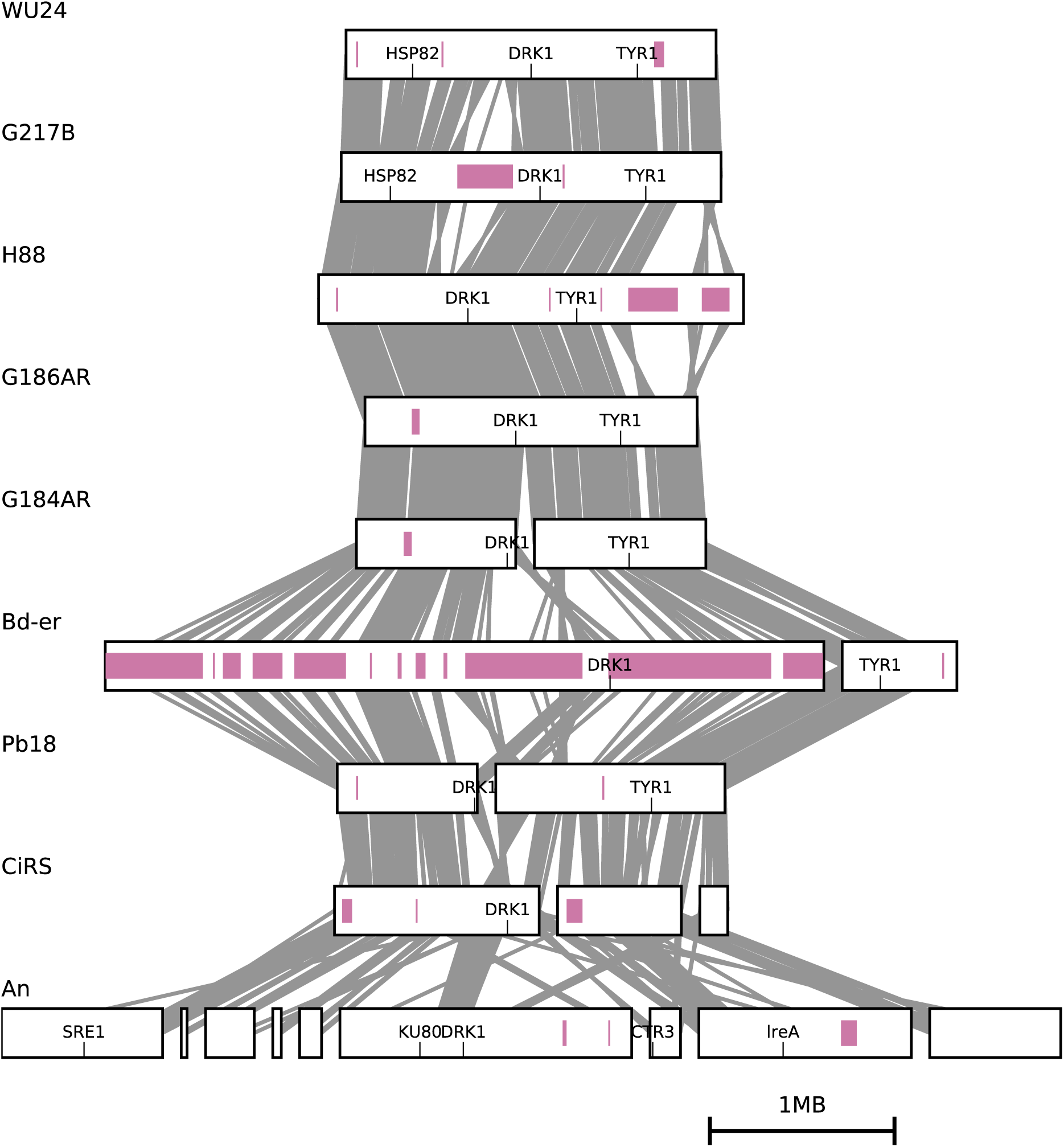
The *DRK1* region is syntenically conserved in *Onygenales*. Chromosome 5 of WU24 aligned to syntenic regions of selected genomes, rendered as in Fig. S9.

**Fig. S12.**
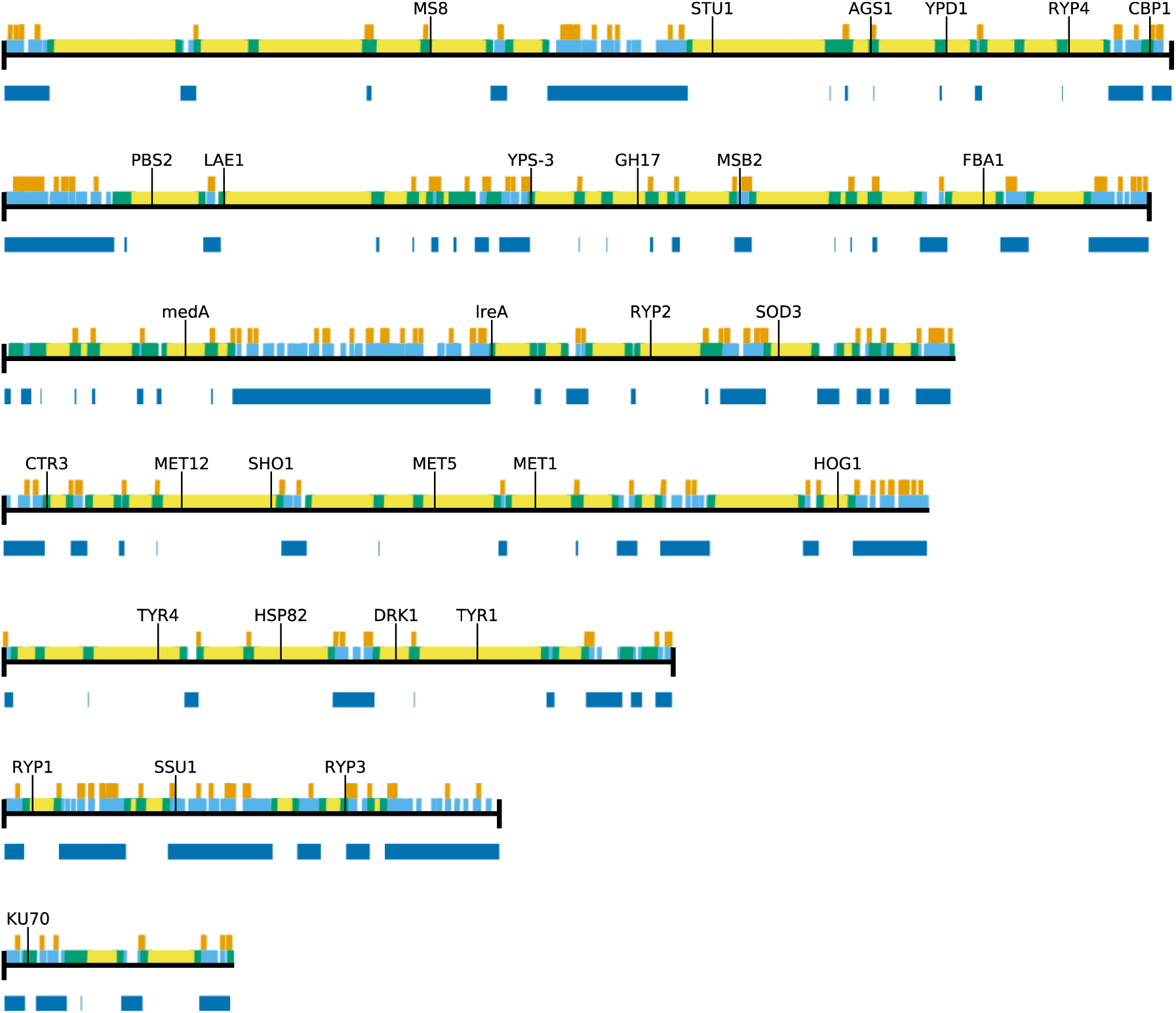
Repetitive regions in G217B. G217B genome assembly with genes colored to show repeat context: light blue for genes in LTR blocks and green for genes near LTR blocks. Transposon genes are shown in orange above the chromosomes and LTR blocks are shown in dark blue below the chromosomes.

**Fig. S13.**
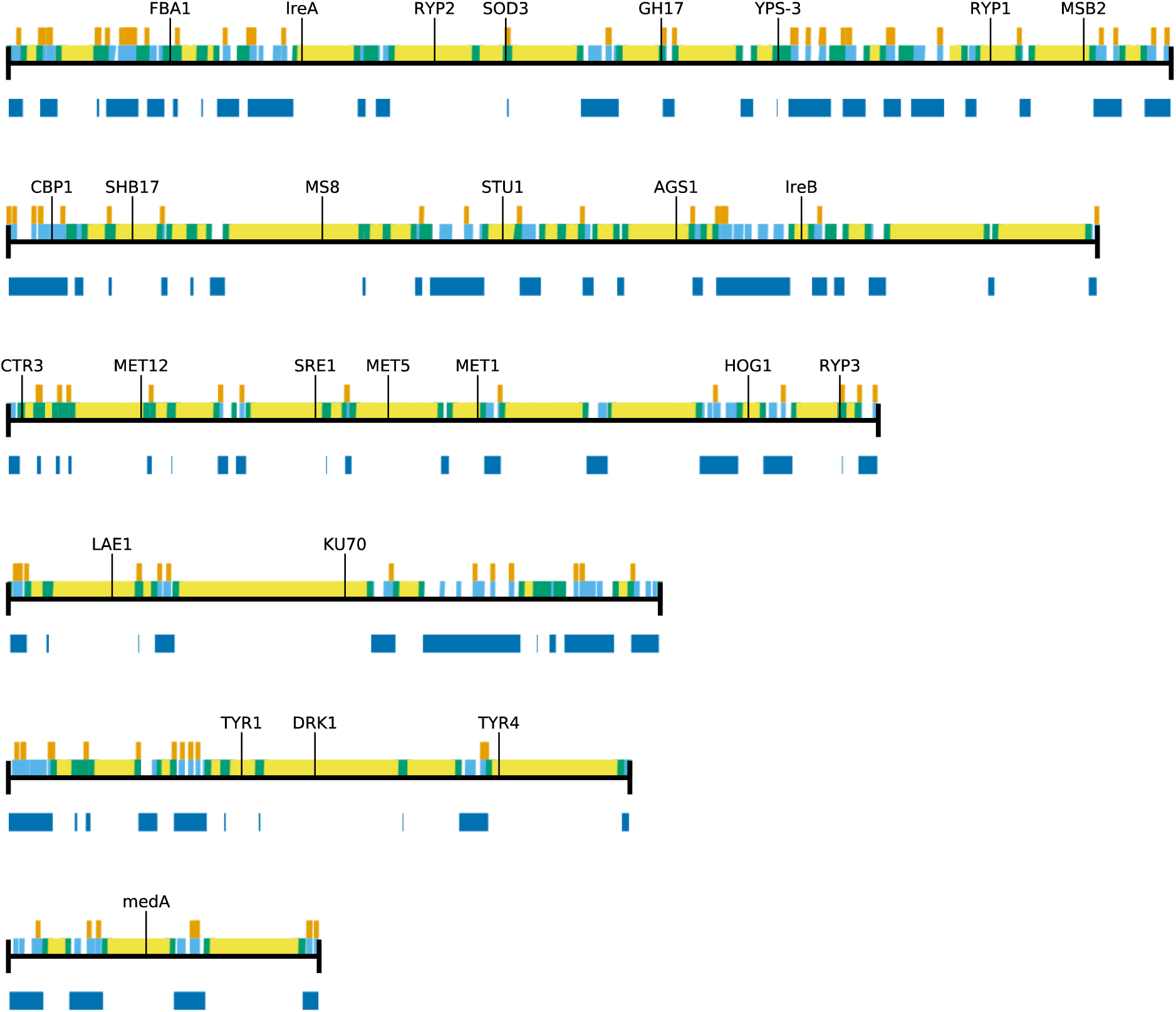
Repetitive regions in H88. H88 genome assembly rendered as in Fig. S12.

**Fig. S14.**
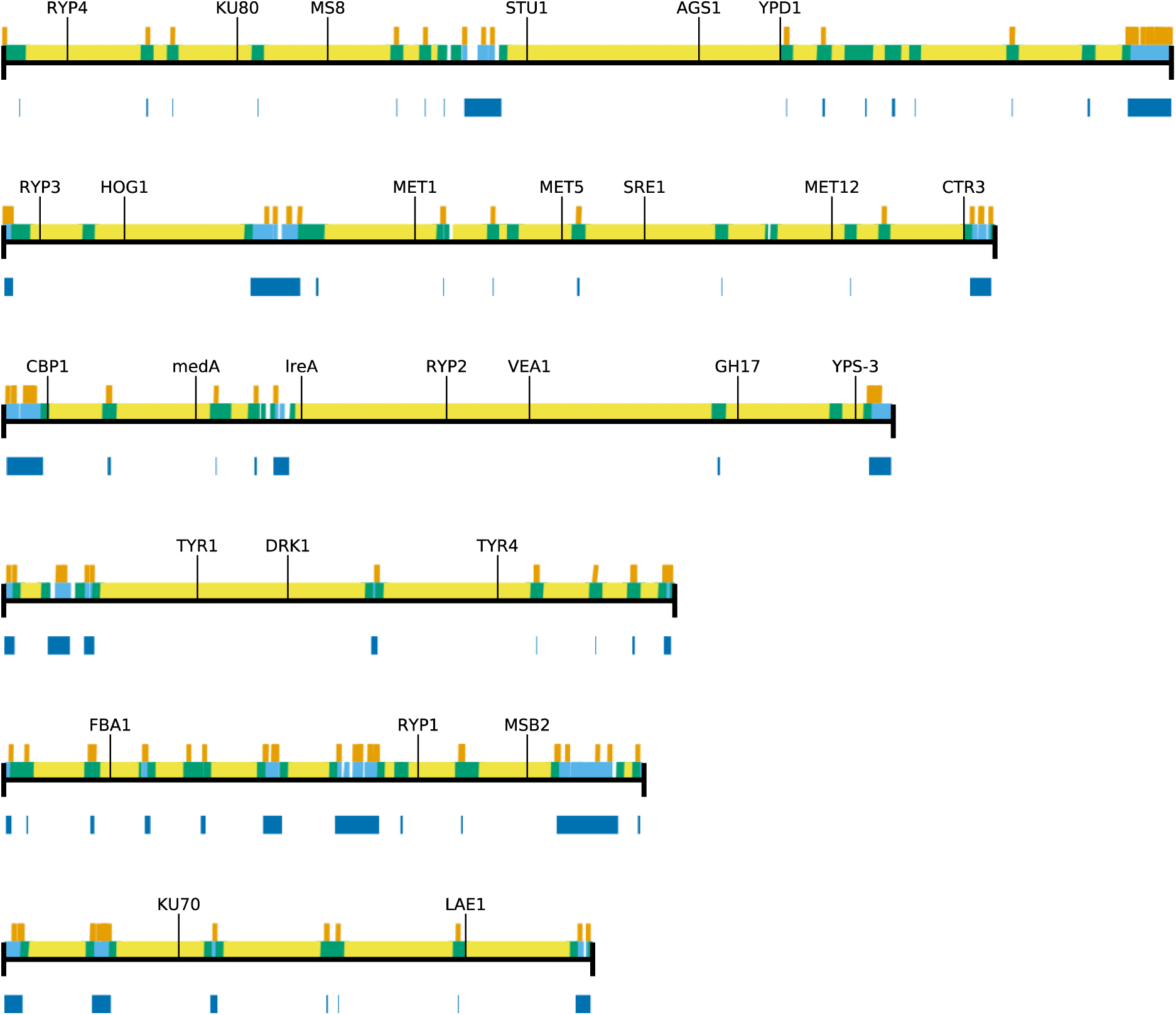
Repetitive regions in G186AR. G186AR genome assembly rendered as in Fig. S12.

**Fig. S15.**
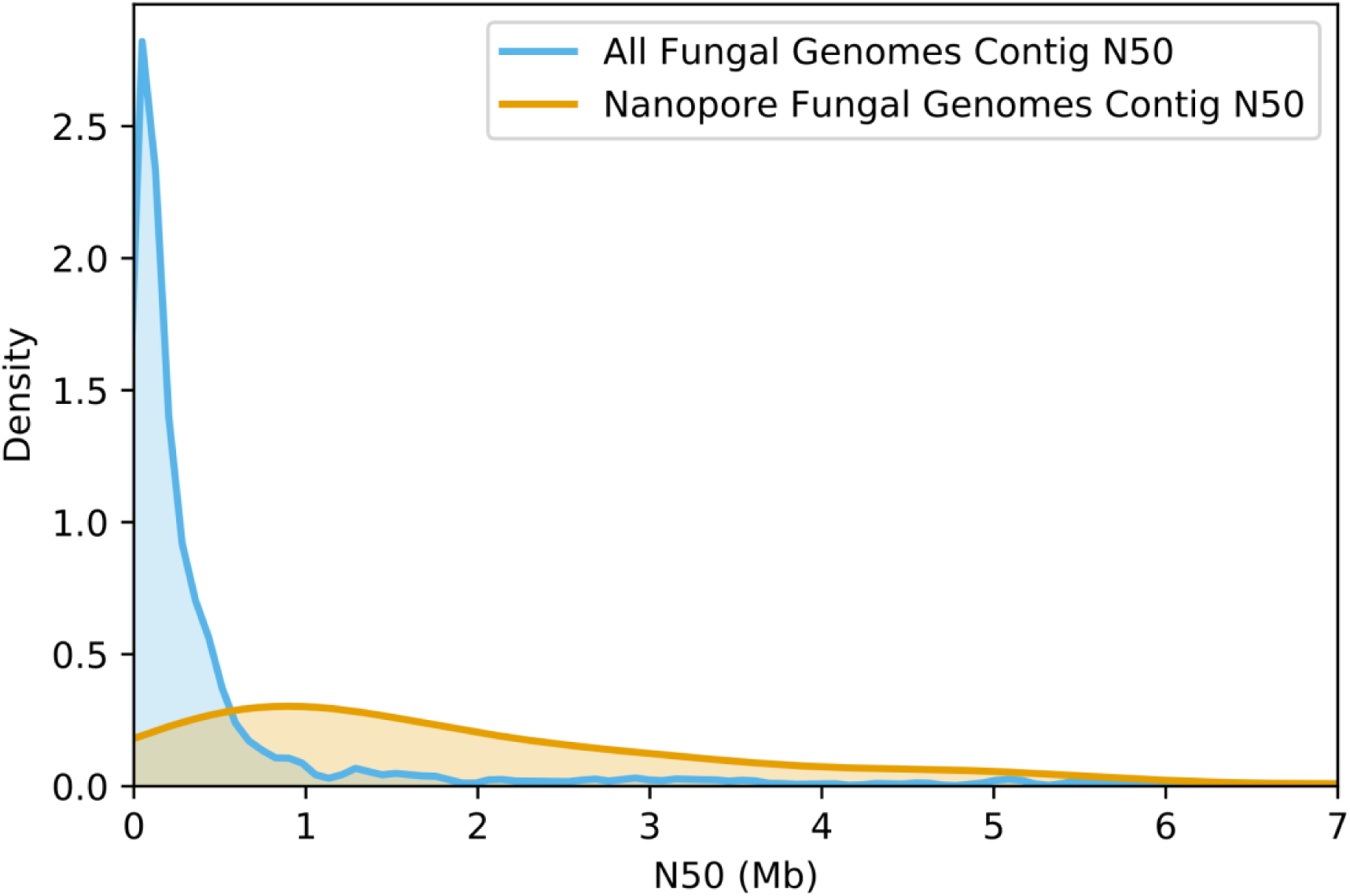
Oxford Nanopore sequencing results in contiguous fungal genomes. Contig N50 values are plotted for 1,395 fungal genomes compared to 137 fungal genomes assembled by Oxford Nanopore reads. Y-values are kernel density estimations of the distributions.

